# Parameter Estimation and Model Selection for the Quantitative Analysis of Oncolytic Virus Therapy in Zebrafish^⋆^

**DOI:** 10.1101/2025.04.23.649270

**Authors:** Yuhong Liu, Dilan Pathirana, Jan Hasenauer

## Abstract

Oncolytic virus therapy (OVT) is emerging as a potent alternative to conventional cancer treatments by employing engineered viruses that selectively infect and lyse tumor cells while sparing normal tissues. Although mathematical models have been developed to elucidate the dynamics of OVT and inform personalized therapies, they are often specific to certain organisms. Mathematical models tailored to more recently developed animal models of OVT, such as zebrafish, are not yet available. Here, we introduce the first mathematical model of OVT trained on zebrafish data from published studies to bridge the gap. We explore a variety of mathematical model structures and perform parameter estimation and model selection. The selected model effectively captures the observed tumor dynamics, i.e. delayed tumor shrinkage, and provides valuable insights into the underlying mechanisms of OVT in zebrafish. Our work establishes the groundwork for advancing experimental studies in zebrafish, contributing to the design of more effective cancer treatment strategies in the future.

## 1. INTRODUCTION

Oncolytic virus therapy (OVT) is a promising alternative to conventional cancer treatments and has demonstrated success in recent human clinical trials (Ling et al., 2023). OVT employs oncolytic viruses (OVs) to eradicate malignant cancer cells through two main mechanisms. Firstly, these viruses are typically engineered strains that selectively infect, replicate within and promote lysis in tumor cells while remaining safe for normal cells (Fig. 1a). Secondly, the lysis of tumor cells by OVs exposes tumor antigens, thereby stimulating the immune system to recognize and attack cancer cells. The latter process helps overcome immunosuppression, a significant challenge since tumor cells often appear non-threatening to the immune system. Antonio Chiocca (2002) and Tian et al. (2022) have comprehensively summarized different types of OVs, how their selectivity is engineered, and how they enhance the effects of antitumor immunity.

**Fig. 1.**
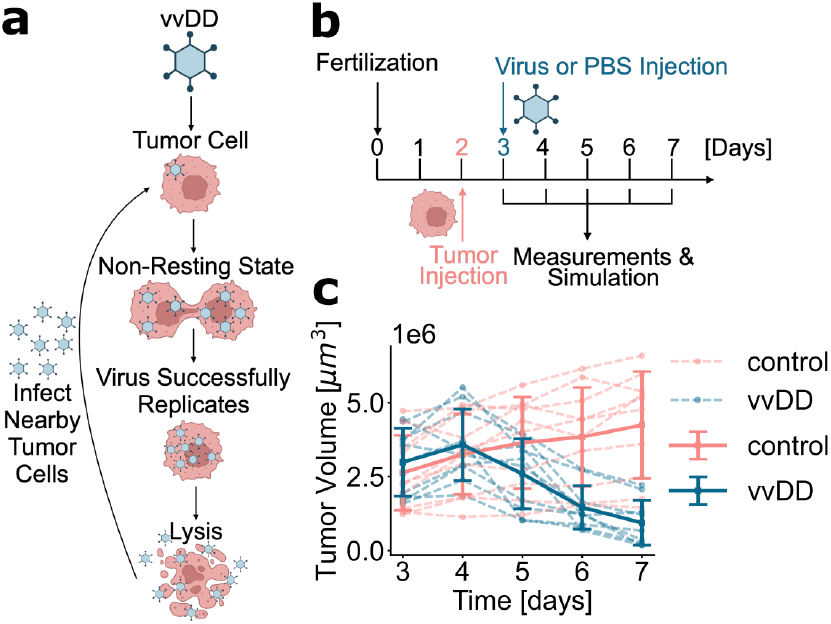
OVT mechanism and zebrafish experiment. **(a)** vvDD replication requires cell division. **(b)** Experimental setup for zebrafish study. Fertilization occurs on day zero. **(c)** Measured tumor volumes for individual zebrafish (dashed) and population mean (solid, error bars are ± 1 std. dev.). *Sub-figures (a) and (b) were created using Biorender*.*com*.

The experimental assessment of (i) the interaction of OVs and tumor cells, and (ii) the contribution of the two mechanisms described above is non-trivial. Accordingly, mathematical models which capture the underlying processes have been developed. These models provide insights into emergent dynamics, guide future experiments, and build the basis for personalized OVT. Various mathematical models have been used to study various process characteristics: Mahasa et al. (2017) studied how infection of normal cells with OVs in the vicinity of tumor cells enhances OVT; Eftimie et al. (2011) explored biological conditions that could lead to the permanent elimination of tumor cells and examined the multistability and instability of the nonlinear dynamics; and Kumar et al. (2021) approached the design of more effective OVs using a fractional-order delay differential equation model that incorporates the viral lytic cycle. Mathematical models, and especially their parameters, are typically tailored to specific animal models. However, there are animal models that are important for cancer research, such as zebrafish that enable rapid screening of OVT candidates (Fazio et al., 2020) and are used for OVT studies (Mealiea et al., 2021), for which no mathematical models are available.

In this manuscript, we propose the first mathematical model for OVT in zebrafish, to our knowledge. To this end, we consider a range of model topologies and perform parameter estimation and model selection using published experimental data from Mealiea et al. (2021). Our selected model can quantitatively capture the measured dynamics and provide insights into underlying mechanisms contributing to, for example, delayed tumor shrinkage. Subsequently, we refine the model to account for the heterogeneity between individual zebrafish through additional parameters for each individual. We show that the individual numbers of initially homed tumor cells can substantially explain the population variability in tumor response. Finally, we discuss important extensions of our models in the direction of mixed-effects and stochastic models for extinction analysis. Our work represents a foundational first step in the mathematical modeling of OVT on zebrafish data. This enables a platform for future research to advance experimental studies in this animal model that can inform the design of effective cancer treatment strategies.

## 2. DATA AND METHODS

### 2.1 Experimental Setup and Measurement

In this manuscript, we analyze and model the OVT data collected by Mealiea et al. (2021) for zebrafish. In their experimental study (Fig. 1b), 20 zebrafish were each injected with 400 fluorescently-labeled tumor cells on day 2 (after fertilization on day 0). The zebrafish were then split into two groups. The treatment group was injected with 10^9^ plaque-forming units of double-deleted vaccinia virus (vvDD) on day 3, while the control group remained untreated and thus injected with only phosphate-buffered saline (PBS). Both groups were observed until day 7.

vvDD is a double-stranded DNA virus with deletions that increase tumor specificity and safety for other cells (McCart et al., 2001), and was reported to only replicate efficiently in tumor cells. In the zebrafish study (Mealiea et al., 2021), the virus is also fluorescently-labeled.

The tumor volume was recorded daily on days 3 to 7 (Fig. 1c). In the vvDD-treated group, the tumor volume declined from day 4 onwards, while it continued to grow in the control group. Overall, both groups show substantial inter-individual heterogeneity in tumor volume.

### 2.2 Mathematical Modeling

We model the interaction between OVs and tumor cells using ordinary differential equation (ODE) models. These ODE systems are of the form:

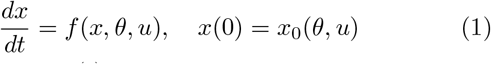

with state vector 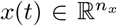 at time *t*, parameter vector 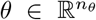, and input vector 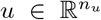 that encodes the experimental conditions. The vector field 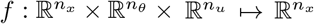 encodes the dynamics of the process and is constructed to be Lipschitz continuous, ensuring that the ODE solution exists and is unique. The function 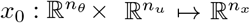 defines the initial condition. The inputs *u* = (*u*_1_, *u*_2_)^*T*^ are used to encode the experimental condition, namely the injected amount of tumor cells (*u*_1_) and virus particles (*u*_2_). The inputs in the vvDD-treated and control conditions are *u*^(*v*)^ = (400, 10^9^)^*T*^ and *u*^(*c*)^ = (400, 0)^*T*^, respectively.

The parameters *θ*, e.g., the growth rate of tumor cells, are inferred from the measured data using maximum likelihood estimation. The observable is the tumor volume,

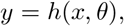

with 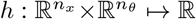 denoting the observation mapping. The zebrafish measurements are denoted by

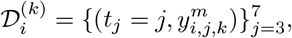

with condition *i* ∈ {*v, c*}, time point index *j* ∈ {3 … 7} in days, and zebrafish individual index *k* ∈ {1 … 10}. The measurement noise is assumed to be additive and normally distributed, *y*^*m*^ = *y* + *ϵ*, with a noise variance depending on the tumor volume, 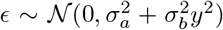, with *σ*_*a*_ *>* 0 and *σ*_*b*_ *>* 0. Accordingly, the conditional probability of observing *y*^*m*^ given *y, σ*_*a*_ and *σ*_*b*_ is given by

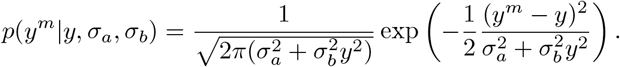

Assuming that the zebrafish are identical, the objective function with negative log-likelihood is

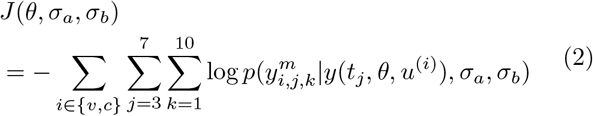

with *y*(*t, θ, u*) = *h*(*x*(*t, θ, u*), *θ*) and *x*(*t, θ, u*) denoting the solution of the ODE model (1) for parameters *θ* and input *u* evaluated at time *t*. To account for differences between zebrafish, we introduce parameters *ξ*_*i*,*k*_ which are specific to the individual zebrafish, yielding the objective function

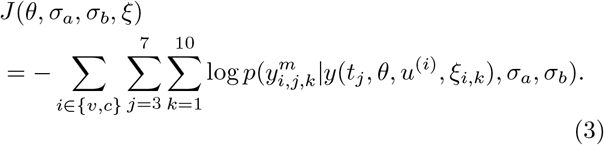

Details on how *ξ* influences the solution are in Section 3.4.

The maximum likelihood estimates are obtained by minimizing the objective functions *J*(*θ, σ*_*a*_, *σ*_*b*_) or *J*(*θ, σ*_*a*_, *σ*_*b*_, *ξ*), i.e., 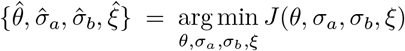 for the latter, subject to parameter bounds, with each parameter bound in ℝ_+_.

### 2.3 Software

We use PEtab (Schmiester et al., 2021) to define the parameter estimation problems and SBML (Hucka et al., 2003) for standardized representations of the mathematical models. Parameter optimization and uncertainty analysis are implemented using pyPESTO (Schälte et al., 2023), wherein we used AMICI (Fröhlich et al., 2021) for simulation and Fides (Fröhlich et al., 2022) for optimization. We used STRIKE-GOLDD to perform structural identifiability analysis (Díaz-Seoane et al., 2023).

## 3. RESULTS

### 3.1 Model formulation and analysis

In the **baseline model**, we distinguish uninfected tumor cells (*U*), infected tumor cells (*I*), and extracellular virus (*V*). The cells are denoted by [*U*], [*I*] and [*V*], yielding the state vector *x* = ([*U*], [*I*], [*V*])^*T*^. This state vector changes over time due to six processes (Fig. 2a):

**Fig. 2.**
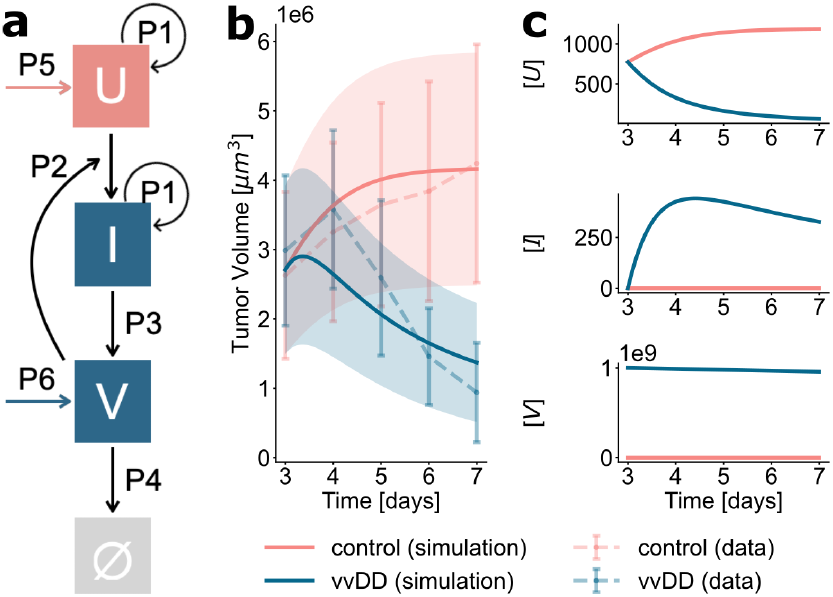
The baseline model. **(a)** Process diagram. **(b)** Optimized model simulation. Data are means with ± 1 empirical std. dev. error bars. Simulations have ± 1 estimated std. dev. error bands. **(c)** Optimized state variable trajectories.

(P1) The quantity of uninfected and infected tumor cells increases due to tumor cell division,

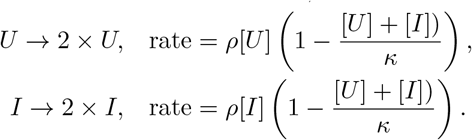

We assume a logistic growth model with maximal division rate constant *ρ* and carrying capacity *κ*.

(P2) The quantity of uninfected tumor cells decreases and the quantity of infected tumor cells increases due to infection with the virus,

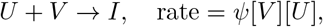

with infection rate constant *ψ*.

(P3) The quantity of infected tumor cells decreases due to cell death, resulting in the release of viruses,

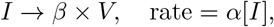

with age-of-infection–independent death rate constant *α*, and the quantity of released virus particles *β*.

(P4) The quantity of extracellular viruses decreases due to virus removal processes,

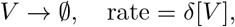

with removal rate constant *δ*.

(P5, P6) Tumor cell injection on day 2 and virus injection on day 3 are processes happening at discrete time points and marking singular events. At these injection points, the respective abundances become nonzero:

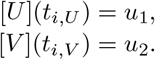

The injection time points are denoted by *t*_*i*,*U*_ = 2 and *t*_*i*,*V*_ = 3 (Fig. 1b), while the quantity of injected uninfected tumor cells and viruses are denoted by *u*_1_ and *u*_2_.

Given the processes P1 to P6, the dynamics can be separated into three intervals:

Interval 1 is the time interval before the injection of tumor cells. In this interval, all state variables are zero:

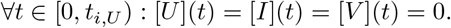

Interval 2 is the time interval between the injections of tumor cells and viruses. In this interval, the quantity of infected tumor cells and viruses is zero, and the quantity of uninfected tumor cells grows (P1):

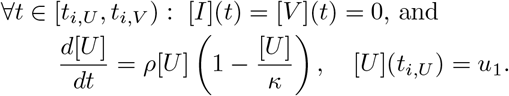

The time-dependent state [*U*](*t*) can be computed analytically using the separation of variables, yielding

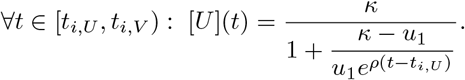

Interval 3 is the time interval after the injection of the virus. In this interval, all state variables can change dynamically:

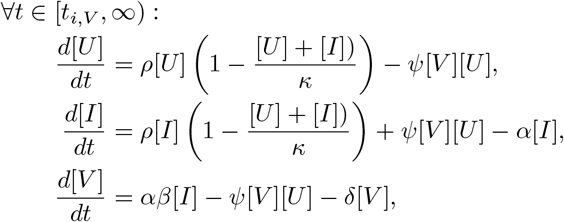

with initial conditions 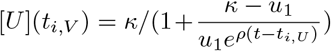 (the final value of the previous interval), [*I*](*t*_*i*,*V*_) = 0, and [*V*](*t*_*i*,*V*_) = *u*_2_. For this interval, no analytical solution is known, and the ODE solution is approximated with AMICI.

The theoretical analysis of the model reveals that it is Lipschitz continuous (proof provided in the Supplementary Material). Furthermore, for non-negative values of the parameters and inputs, the non-negative quadrant, i.e. 0 ≤ [*U*], 0 ≤ [*I*], 0 ≤ [*V*], is invariant. This means that, with an initial condition in this quadrant, the states remain non-negative, which is biologically reasonable.

The data provide information about the volume of the tumor, which is assumed to be proportional to the quantity of tumor cells. Thus, the observable mapping is

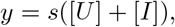

where the scaling factor *s* is the mean volume of a tumor cell and y is the tumor volume. Structural identifiability analysis with STRIKE-GOLDD reveals that this parameter is non-identifiable (see Supplementary Material). To resolve this, we searched the literature to approximate *s*. The tumor cells used in the experiment are from the MC38 murine colorectal adenocarcinoma cell line, which has an average doubling time of about 24 hours (Morimoto-Tomita et al., 2005). Accordingly, one would expect 800 tumor cells at day 3. As the mean tumor volume at day 3 is 2 808 614 µm^3^, we estimate 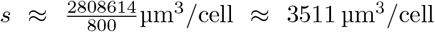. With this estimate of *s*, the baseline model achieved “Full Input-State-Parameter Observability” (FISPO) (Díaz-Seoane et al., 2023), suggesting that with sufficient measurement data of *y*, all parameters are identifiable. A limitation of this approximation is that the tumor doubling time during the first day is assumed to be comparable between zebrafish and mice.

### 3.2 Parametrization of Baseline Model

To assess whether the baseline model can describe the data, we obtain the maximum likelihood estimate by minimizing (2). Individual measurements are used for the optimization but the heterogeneity between individual zebrafish is not accounted for in the baseline model. Parameter bounds are provided in Table 1.

**Table 1.** Informed parameter bounds of all models. The lower and upper bounds for the parameters and the reasoning are provided. For parameters that are not defined, a lower bound of 1e-8 and an upper bound of 1e8 is used.

| Parameter | Effect | Bounds | Information |
| --- | --- | --- | --- |
| $\rho$ | Cell division | [0.42, 1.66] | Tumor doubling time is $\approx 1$ day (Morimoto-Tomita et al., 2005) |
| $\kappa$ | Carrying capacity | [1e2, 1e5] | Tumor growth slows at day 7 with $< 2000$ tumor cells (scaled by $s$) (Fig. 1) |
| $\psi$ | Infection | [1e-10, 1e-2] | Arbitrary interval based on estimate from Iwami et al. (2012) |
| $\beta$ | Virus burst size | [1e0, 1e4] | Arbitrary interval based on estimate from Iwami et al. (2012) |
| $\alpha$ | Cell death | [1e-4, 1e3] | Arbitrary interval based on estimate from Iwami et al. (2012) |
| $\delta$ | Virus degradation | [1e-2, 1e2] | Arbitrary interval based on estimate from Neumann et al. (1998) |
| $\xi_{i,k}$ | Individual homing | [1e-7, 1e0] | Scaling factor for effective tumor inoculation amount, necessarily $< 1$ |
| $\sigma_a$ | Additive noise | [1e1, 1e5] | Similar upper bound as $\kappa$ |
| $\sigma_b$ | Proportional noise | [1e-4, 1e2] | Pairwise relative differences in the data were always $< 20$ (Fig. 1) |

**Table 2.** AIC and AICc values for different model candidates.

| Model | AIC | AICc |
| --- | --- | --- |
| Baseline | 1475.8 | 1477.3 |
| Age-of-infection | 1464.7 | 1466.7 |
| Individual-based | **1437.0** | **1461.8** |

The analysis of the fitting results reveals that the optimization converges robustly, suggesting that globally optimal parameters are found (see Supplementary Material). The model provides a good description of the measurement data (Fig. 2b). However, it fails to capture the quantitative dynamics of the tumor volume after virus injection on day 3. Although the measurements show that mean tumor volume increases until day 4, the best model fit exhibits a decrease. Hence, the baseline model cannot capture the observed delay in tumor shrinkage.

To assess parameter uncertainties, we compute 95% confidence intervals with the profile likelihood method (Raue et al., 2009). This reveals that the parameters *α, κ, ψ*, and *σ*_*b*_ are practically identifiable, as their confidence intervals are finite (Fig. 3). The parameters *δ, ρ, σ*_*a*_, and *β* cannot be bounded from both above and below, and hence are practically non-identifiable.

**Fig. 3.**
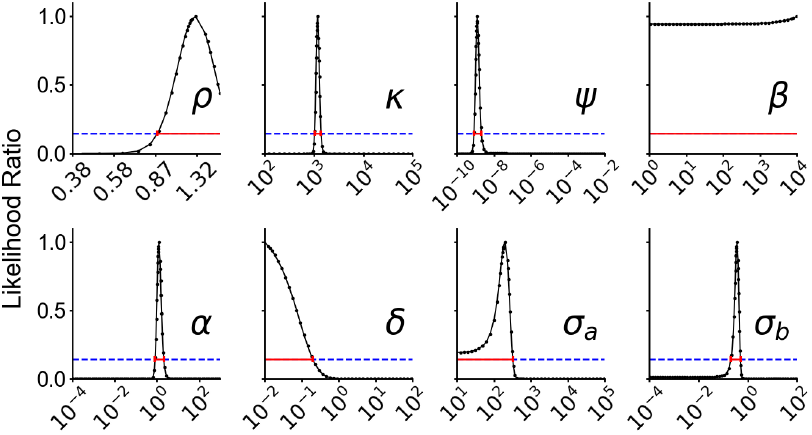
Baseline model profile likelihoods. Domains are the parameter bounds (Table 1). Blue lines are the likelihood ratio threshold that was used to compute the 95% confidence intervals (red lines).

### 3.3 Modeling delayed tumor shrinkage

As the baseline model fails to describe key properties of the measurement data, we consider an **age-of-infection model**, which accounts for the experimental observation that cell lysis does not occur directly after infection but instead after a large number of virus particles are produced (Dominguez et al., 2015). This could be captured using a discrete delay (and a delay differential equation), yet, here we use the linear chain trick (Hurtado et al., 2020). Therefore, we split the state variable for infected cells *I* into *L* state variables, *I*_*l*_, with *l* ∈ {1 … *L*}. Tumor cells in state *I*_1_ are infected but carry a low viral load, while tumor cells in state *I*_*L*_ carry a high viral load. Cell division occurs for all tumor cells at the same rate (P1), but the infection of uninfected tumor cells only yields *I*_1_ (P2). Infected cells in state *I*_*l*_ (except *l* = *L*) can transition into state *I*_*l*+1_ according to process P2’:

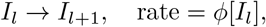

which captures different stages of the infected cell due to an increase in the number of virus particles, with the transition rate constant *ϕ*. Here we assume one constant transition parameter for simplicity. Cells that reach the state *I*_*L*_ can undergo cell lysis, which results in virus release (P3). This yields the governing ODE system for interval 3, ∀*t* ∈ [*t*_*i*,*V*_, ∞):

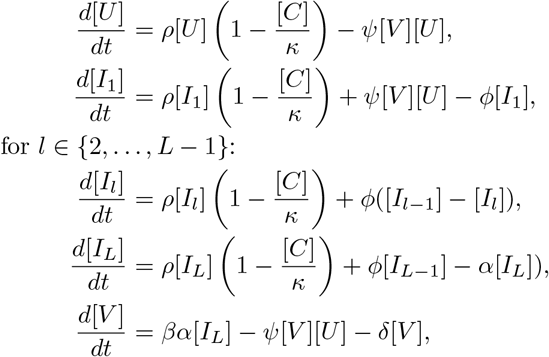

with 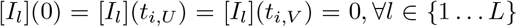, the same initial conditions as in the baseline model for [*U*] and [*V*], and total tumor cell number 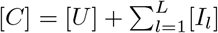. In this study, we use *L* = 5.

We estimate the parameters of the age-of-infection model using the same approach as for the baseline model, with the corresponding observable mapping *y* = *s*[*C*]. The optimization also achieved good convergence, suggesting that the global optimum is found (see Supplementary Material). The assessment of the fit reveals a good agreement with the measurement data (Fig. 4b). Indeed, the model simulation exhibits a delay in tumor shrinkage. The obtained state variables’ trajectories show that virus accumulation (Fig. 4c) causes a delay.

**Fig. 4.**
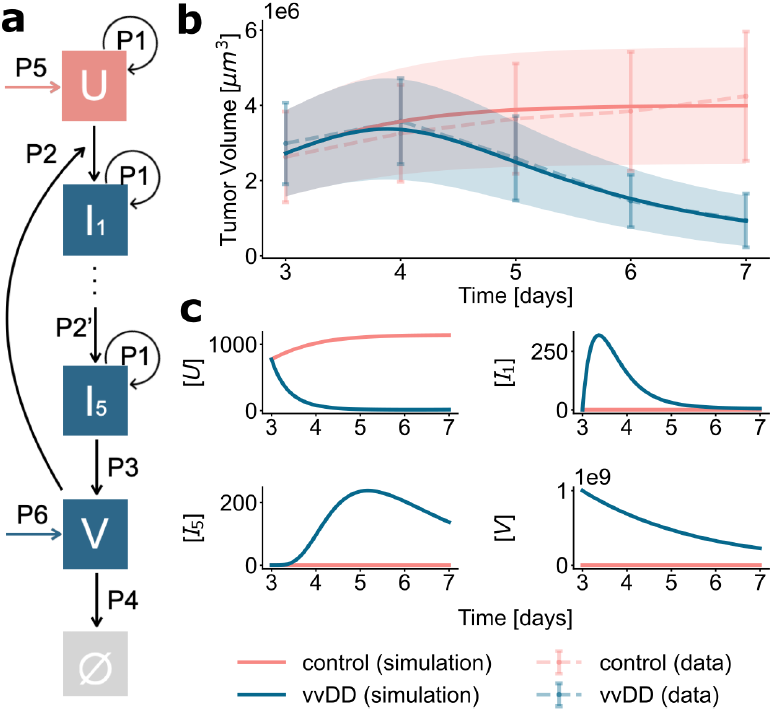
The age-of-infection model. **(a)** Process diagram. **(b)** Optimized model simulation. Data are means with ±1 empirical std. dev. error bars. Simulations have ± 1 estimated std. dev. error bands. **(c)** Optimized state variable trajectories.

We used the same literature-based estimate of *s* as in the baseline model. The structural identifiability analysis with STRIKE-GOLDD found that the model is FISPO. Assessment of the parameter uncertainties revealed that *ρ, ψ, β*, and *δ* are practically non-identifiable (Fig. 5).

**Fig. 5.**
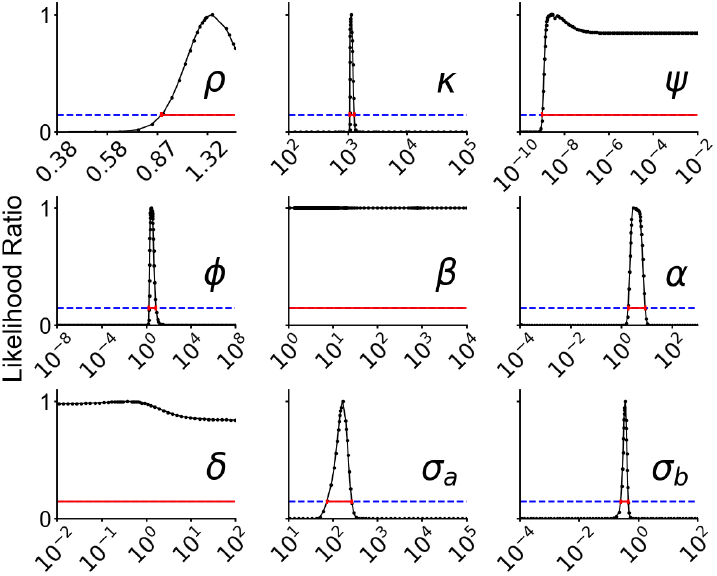
Age-of-infection model profile likelihoods. Domains are the parameter bounds (Table 1). Blue lines are the likelihood ratio threshold that was used to compute the 95% confidence intervals (red lines).

### 3.4 Modeling Inter-Individual Variability

The age-of-infection model provides a good description of the mean tumor growth and shrinkage dynamics amongst zebrafish. Yet, it does not explain the observed inter-individual variability. We thus consider an **individual-based** age-of-infection model; specifically, we allow for differences in the initial amount of tumor cells, e.g., due to different numbers of injected cells or different homing rates. Accordingly, the initial condition in time interval 2 can differ between individual zebrafish,

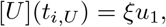

with individual-specific homing parameter 0 *< ξ* ≤ 1.

We estimate the shared and individual parameters of the model by minimizing objective function (3). In contrast to the previous estimation problem for the age-of-infection model, we have 20 additional parameters: *ξ*_*v*,1_ to *ξ*_*c*,10_. Despite this increase, we observe that the optimization also converges (see Supplementary Material).

The assessment of the fitting results reveals relatively good agreement of data and model simulation for individual zebrafish (Fig. 6). Indeed, individual-specific homing parameters describe individual dynamics better for the most part. However, not all variance is explained, as shown in some individuals. Model selection using the standard (AIC) and the corrected (AICc) Akaike Information Criterion shows that the individual-based age-of-infection model performs best (Tab. 2). Accordingly, the individual-based model provides a better balance between model fit and model complexity. The uncertainty analysis using the profile likelihood reveals that some parameters are practically non-identifiable (see Supplementary Material).

**Fig. 6.**
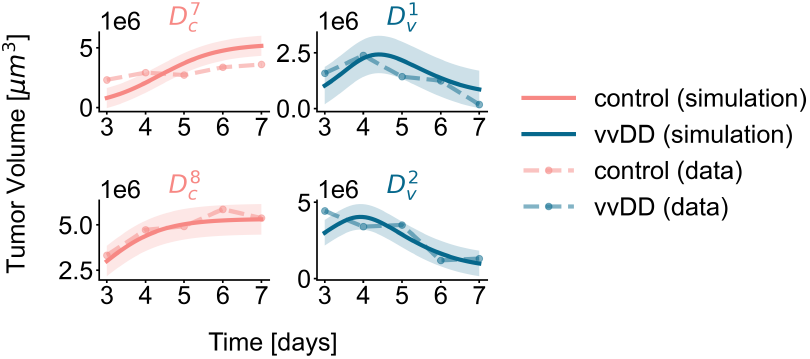
Optimized fits with the individual-based model. Simulations have ±1 estimated std. dev. error bands. The fits for the remaining 16 zebrafish are provided in the Supplementary Material.

## 4. DISCUSSION

OVT is a promising alternative to established anti-tumor therapies. Yet, the understanding of the underlying processes is limited. Here, we presented three mathematical models for OVT in zebrafish, an experimental animal model that enables rapid screening of OVT candidates. Our analysis reveals that a baseline model cannot capture the delay in tumor shrinkage. In contrast, the observed delay in tumor volume reduction can be accurately described using the age-of-infection model with multiple infection states representing different intracellular virus concentrations. The formulation of the individual-based age-of-infection model, which accounts for differences in the number of homed tumor cells, provides a better description of the observed variability.

Yet, our analysis also reveals that the homing rates of tumor cells do not fully capture the observed variability. As a next step, we consider two model extensions that may improve this. Firstly, the joint fitting of the individuals could be improved by with nonlinear mixed-effects models. Such models allow for improved integration and dissection of inter-individual variability and uncertainty. This can be particularly beneficial in the case of non-identifiability. Besides *ξ*, additional inter-individual differences, e.g., individual-specific carrying capacity *κ*, could also be considered. A structured approach should then be used to select the most relevant sources of interindividual variability. Secondly, tumor clearance is inherently stochastic due to heterogeneity at low tumor cell numbers. An immediate extension of our study is to perform extinction analysis with a stochastic implementation of the described biological processes. This would enable identification of OVT parameters that can optimally drive tumors to extinction.

By making our code and models publicly available, we aim to facilitate reuse of our work and support further research in this area. The proposed models provide a foundation for future studies to advance OVT strategies.

## Supporting information

All additional figures and results, together with all SBML and PEtab files, and the code and instructions to perform all analyses

## ACKNOWLEDGEMENTS

We thank the members of the research group of Prof. Dr. Jan Hasenauer for valuable feedback on drafts. We acknowledge the Marvin and Unicorn clusters hosted by the University of Bonn.

## SUPPLEMENTARY MATERIAL

All additional figures and results, together with all SBML and PEtab files, and the code and instructions to perform all analyses, is available at https://doi.org/10.5281/zenodo.14802838.

