## Supplementary material for "Parameter Estimation and Model Selection for the Quantitative Analysis of Oncolytic Virus Therapy in Zebrafish^⋆^": All additional figures and results, together with all SBML and PEtab files, and the code and instructions to perform all analyses: check_gradients.ipynb

```
# %% [code]
import numpy as np
import pandas as pd
import seaborn as sns

import petab
import pypesto
import pypesto.petab


# %% [code]
# rng seed
np.random.seed(500)

# %% [markdown]
Import petab files and define the problem

# %% [code]
petab_yaml = 'petab_files/age_of_infection_model.yaml'
petab.validate(petab_yaml)
petab_problem = petab.Problem.from_yaml(petab_yaml)

problem = pypesto.petab.PetabImporter(
        petab_problem,
        hierarchical=False,
        model_name=f"Age_of_Infection_Model",
    ).create_problem(force_compile=True, verbose=False)
problem.objective.amici_model.setAllStatesNonNegative()

# %% [markdown]
# Gradient Check

# %% [code]
# obtain random startpoints
startpoints = problem.get_startpoints(n_starts=2)

# %% [code]
problem.objective.check_grad(
    x = startpoints[0],
    eps = 1e-5,  # default
    verbosity = 0,
)

# %% [code]
problem.objective.check_grad(
    x = problem.get_reduced_vector(startpoints[0]),
    eps = 1e-5,  # default
    verbosity = 0,
)

# %% [code]
gc = problem.objective.check_grad_multi_eps(
    x=startpoints[0],
    verbosity=0,
    label='rel_err',  # default
)
gc

# %% [code]
def highlight_value_above_threshold(x, threshold=10):
    return ['color: darkorange' if xi > threshold else None for xi in x]

gc.style.apply(
    highlight_value_above_threshold, subset=["fd_err"],
).background_gradient(
    cmap=sns.light_palette("purple", as_cmap=True), subset=["abs_err"],
).background_gradient(
    cmap=sns.light_palette("red", as_cmap=True), subset=["rel_err"],
)

# %% [code]
```
