## Supplementary material for "Parameter Estimation and Model Selection for the Quantitative Analysis of Oncolytic Virus Therapy in Zebrafish^⋆^": All additional figures and results, together with all SBML and PEtab files, and the code and instructions to perform all analyses: model_creation.py

```
import tellurium as te
import libsbml

# Generate ODE functions and initial conditions
ode_functions = []
initial_conditions = []
L = 5

# ODE for U
ode_functions.append("U' = rho * U * (1 - (U + " + " + ".join([f"I_{j}" for j in range(1, L+1)]) + ")/kappa) - psi * V * U;")
initial_conditions.append("U = kappa / (1 + (kappa/400 - 1) * exp(-rho));")

# ODE for I_1
ode_functions.append("I_1' = rho * I_1 * (1 - (U + " + " + ".join([f"I_{j}" for j in range(1, L+1)]) + ")/kappa) + psi * V * U - phi * I_1;")
initial_conditions.append("I_1 = 0;")

# ODE and initial conditions for I_2 to I_{L-1}
for j in range(2, L):
  ode_functions.append(f"I_{j}' = rho * I_{j} * (1 - (U + " + " + ".join([f"I_{k}" for k in range(1, L+1)]) + f")/kappa) + phi * I_{j-1} - phi * I_{j};")
  initial_conditions.append(f"I_{j} = 0;")

# ODE for I_L
ode_functions.append(f"I_{L}' = rho * I_{L} * (1 - (U + " + " + ".join([f"I_{j}" for j in range(1, L+1)]) + f")/kappa) + phi * I_{L-1} - alpha * I_{L};")
initial_conditions.append(f"I_{L} = 0;")

# ODE for V
ode_functions.append(f"V' = beta * alpha * I_{L} - psi * V * U - delta * V;")
initial_conditions.append("V = u_2;")

age_of_infection_model = f"""
model age_of_infection_model

  // ODE functions
  {' '.join(ode_functions)}

  // initial conditions
  // t=0 is when the virus is injected
  {' '.join(initial_conditions)}

  // condition dependent parameters
  u_2 = 1 * 1E9; // pfu
  
  // parameters to be estimated
  rho = 0.01;
  kappa = 0.01;
  psi = 0.01;
  phi = 0.01;
  alpha = 0.01;
  beta = 0.01;
  delta = 0.01;

  // Event to set U to zero when U falls below 1e-8, to avoid exploding solutions that are not biologically relevant
  at (U < 1e-8): U = 0;
end
"""

# Load the model
r = te.loada(age_of_infection_model)
# Export the model to SBML
r.exportToSBML('petab_files/age_of_infection_model.xml', current=False)
```
