## Supplementary material for "Parameter Estimation and Model Selection for the Quantitative Analysis of Oncolytic Virus Therapy in Zebrafish^⋆^": All additional figures and results, together with all SBML and PEtab files, and the code and instructions to perform all analyses: model_optimization.py

```
import os
import numpy as np
import petab
import pypesto
import pypesto.petab
from pypesto.optimize import minimize, FidesOptimizer
from pypesto.engine import MultiProcessEngine

def main():    

    ###########################################################################
    ################ set the number of starts and maxiter #####################
    n_start, maxiter = 5000, 5000
    ###########################################################################


    # set the number of processes
    n_procs = int(os.environ.get("SLURM_CPUS_ON_NODE", os.cpu_count()))

    # load the petab problem
    petab_yaml = 'petab_files/age_of_infection_model.yaml'
    petab.validate(petab_yaml)
    petab_problem = petab.Problem.from_yaml(petab_yaml)

    np.random.seed(500)

    problem = pypesto.petab.PetabImporter(
            petab_problem,
            hierarchical=False,
            model_name=f"Age_of_Infection_Model",
        ).create_problem(force_compile=True)
    problem.objective.amici_model.setAllStatesNonNegative()

    # run the optimization
    result = minimize(
        problem=problem,
        optimizer=FidesOptimizer(verbose=False, options={'maxiter': maxiter}),
        n_starts=n_start,
        engine=MultiProcessEngine(n_procs=n_procs),
        history_options = pypesto.HistoryOptions(trace_record=False, storage_file='optimization_history/age_of_infection_model.hdf5'),
        filename='optimization_history/age_of_infection_model.hdf5',
    )

    # print result summary
    print(result.summary())

if __name__ == "__main__":
    main()
```
