## Supplementary material for "Parameter Estimation and Model Selection for the Quantitative Analysis of Oncolytic Virus Therapy in Zebrafish^⋆^": All additional figures and results, together with all SBML and PEtab files, and the code and instructions to perform all analyses: age_of_infection_model.yaml

```
format_version: 1
parameter_file: parameters.tsv
problems:
  - condition_files:
    - conditions.tsv
    measurement_files:
    - measurements_scaled.tsv
    observable_files:
    - observables.tsv
    sbml_files:
    - age_of_infection_model.xml
    visualization_files:
    - visualizations.tsv
```
