## Supplementary material for "Parameter Estimation and Model Selection for the Quantitative Analysis of Oncolytic Virus Therapy in Zebrafish^⋆^": All additional figures and results, together with all SBML and PEtab files, and the code and instructions to perform all analyses: petab_files_creation.ipynb

```
# %% [code]
import pandas as pd
import numpy as np
import matplotlib.pyplot as plt
import os

# %% [code]
folder_path = '../../data/'

# Plot setting
plt.rcParams['font.size'] = 30

dpi = 100
wid = int(8)
hei = int(6)

red_color = '#f78884'
blue_color = '#06688c'

# %% [markdown]
# Obtain data from data.csv

# %% [code]
"""
first, we need to import all the data in data/data.csv
which has gathered all tumor volume data in McCart 2021 et al. with webplotdigitizer

data structure:
1. the first 20 columns are the tumor volume under vvDD condition
   specifying x (time) and y (tumor volume) for each two columns, totalling 10 samples
   the first row can be ignored
   the 1-5 rows are the 5 time points for all 10 samples
2. the next 20 columns are the tumor volume under control/PBS condition with similar structure
3. the last four columns are the mean and stde data of the tumor volume under vvDD condition and control/PBS condition
   each two columns specify one condition, respectively the x and y
   each three rows specify one time point: mean, stde+, and stde-
   again, the first row can be ignored

"""

# Read the CSV file
df = pd.read_csv(os.path.join(folder_path, 'data.csv'))

# Extract tumor volume data for vvDD condition
# Select every other column starting from the second column (Y values)
tumor_vol_vvDD = df.iloc[1:6, 1:20:2].values  
# Create a DataFrame with known time points and extracted tumor volumes 
tumor_vol_vvDD_df = pd.DataFrame(tumor_vol_vvDD, 
                                 index=[0, 1, 2, 3, 4], # ([0,1,2,3,4] is used for time instead of [3,4,5,6,7])
                                 columns=[f'vvDD_sample_{i+1}' for i in range(10)])

# Extract tumor volume data for control/PBS condition
# Select every other column starting from the second column (Y values)
tumor_vol_pbs = df.iloc[1:6, 21:40:2].values
# Create a DataFrame with known time points and extracted tumor volumes
tumor_vol_pbs_df = pd.DataFrame(tumor_vol_pbs, 
                                index=[0, 1, 2, 3, 4], # ([0,1,2,3,4] is used for time instead of [3,4,5,6,7])
                                columns=[f'pbs_sample_{i+1}' for i in range(10)])

# Extract mean and standard deviation data for vvDD and PBS conditions, to verify the data above later
vvDD_mean_stde_data = df.iloc[1:, -3].values
pbs_mean_stde_data = df.iloc[1:, -1].values
# Create DataFrames for vvDD and PBS data
vvDD_mean_stde_df = pd.DataFrame(vvDD_mean_stde_data.reshape(-1, 3), 
                                 index=[0, 1, 2, 3, 4], 
                                 columns=['mean', 'stde_plus', 'stde_minus'])

pbs_mean_stde_df = pd.DataFrame(pbs_mean_stde_data.reshape(-1, 3), 
                                index=[0, 1, 2, 3, 4], 
                                columns=['mean', 'stde_plus', 'stde_minus'])

# convert data to numeric type
tumor_vol_vvDD_df = tumor_vol_vvDD_df.apply(pd.to_numeric, errors='coerce')
tumor_vol_pbs_df = tumor_vol_pbs_df.apply(pd.to_numeric, errors='coerce')
vvDD_mean_stde_df = vvDD_mean_stde_df.apply(pd.to_numeric, errors='coerce')
pbs_mean_stde_df = pbs_mean_stde_df.apply(pd.to_numeric, errors='coerce')

# %% [markdown]
# Verify data

# %% [code]
# plot the individual tumor volume data for vvDD and PBS condition
# this figure can then be checked with the original data in the paper
plt.figure(figsize=(wid, hei))

# Plot vvDD data
for i in range(10):
    plt.plot(tumor_vol_vvDD_df.index, tumor_vol_vvDD_df.iloc[:, i], 
             marker='o', linestyle='-', lw=3, color=blue_color, alpha=0.4, label='vvDD' if i == 0 else "")

# Plot PBS data
for i in range(10):
    plt.plot(tumor_vol_pbs_df.index, tumor_vol_pbs_df.iloc[:, i], 
             marker='o', linestyle='-', lw=3, color=red_color, alpha=0.4, label='PBS' if i == 0 else "")

plt.xlabel('Time (days)')
plt.ylabel(r'Tumor Volume ($\mu m^3$)')
plt.title('Tumor Volume vs. Time\n(vvDD and PBS Conditions)')
plt.legend()
plt.grid(False)
plt.xticks(np.arange(5), np.arange(5)+3)
plt.legend(loc='lower center', ncol=2, frameon=False, bbox_to_anchor=(0.5, -0.5))
plt.show()

# %% [code]
# plot the mean and stde data for vvDD and PBS condition
# the calculated mean and stde data can be compared with the provided data to verify everything is okay

# Calculate mean and standard error for vvDD
vvDD_mean = tumor_vol_vvDD_df.mean(axis=1)
vvDD_se = tumor_vol_vvDD_df.sem(axis=1)
# Calculate mean and standard error for PBS
pbs_mean = tumor_vol_pbs_df.mean(axis=1)
pbs_se = tumor_vol_pbs_df.sem(axis=1)

# Plot both provided and calculated data
plt.figure(figsize=(wid, hei))
# Plot provided data
plt.errorbar(vvDD_mean_stde_df.index, vvDD_mean_stde_df['mean'], 
             yerr=[vvDD_mean_stde_df['mean'] - vvDD_mean_stde_df['stde_minus'], 
                   vvDD_mean_stde_df['stde_plus'] - vvDD_mean_stde_df['mean']],
             fmt='o-', lw=3, color=blue_color, ecolor='lightblue', capsize=5, capthick=3, label='vvDD (provided)')
plt.errorbar(pbs_mean_stde_df.index, pbs_mean_stde_df['mean'], 
             yerr=[pbs_mean_stde_df['mean'] - pbs_mean_stde_df['stde_minus'], 
                   pbs_mean_stde_df['stde_plus'] - pbs_mean_stde_df['mean']],
             fmt='o-', lw=3, color=red_color, ecolor='lightcoral', capsize=5, capthick=3, label='PBS (provided)')
# Plot calculated data
plt.errorbar(vvDD_mean.index, vvDD_mean, yerr=vvDD_se,
             fmt='s--', lw=3, color='darkblue', ecolor=blue_color, capsize=5,capthick=3,  label='vvDD (calculated)')
plt.errorbar(pbs_mean.index, pbs_mean, yerr=pbs_se,
             fmt='s--', lw=3, color='darkred', ecolor=red_color, capsize=5, capthick=3, label='PBS (calculated)')
plt.xlabel('Time (days)')
plt.ylabel(r'Tumor Volume ($\mu m^3$)')
# plt.title('Verify Data Processing\n(Compare Provided and Calculated mean and std)')
plt.legend()
plt.grid(False)
plt.xticks([0,1,2,3,4], [3, 4, 5, 6, 7])
plt.legend(loc='lower center', ncol=1, frameon=False, bbox_to_anchor=(0.5, -0.9))
plt.show()

# %% [markdown]
*One can check whether all the means and stds are matching with the mean and std obatained directly from the paper, to verify whether the processing of the data is correct*

# %% [markdown]
# Scaling of the data

# %% [code]
# Combine day 3 data for vvDD and PBS samples
day3_vvDD = tumor_vol_vvDD_df.iloc[0]
day3_pbs = tumor_vol_pbs_df.iloc[0]
day3_combined = pd.concat([day3_vvDD, day3_pbs])

# Calculate the average volume at day 3 for all samples
day3_avg = day3_combined.mean()

# Calculate the scaling factor s
s = day3_avg / 800

print(f"Average volume at day 3 for all samples: {day3_avg}")
print(f"Assuming 800 cells at day 3, the average s (scaling factor): {s}")

# %% [code]
# rescale the tumor volume to tumor cell number
scaled_tumor_vol_vvDD_df = tumor_vol_vvDD_df / s
scaled_tumor_vol_pbs_df = tumor_vol_pbs_df / s

# %% [markdown]
# Coefficient of Variation calculation to decide the noise model

# %% [code]
# Calculate the coefficient of variation for vvDD condition
vvDD_cv = scaled_tumor_vol_vvDD_df.std(axis=1) / scaled_tumor_vol_vvDD_df.mean(axis=1)

# Calculate the coefficient of variation for PBS condition
pbs_cv = scaled_tumor_vol_pbs_df.std(axis=1) / scaled_tumor_vol_pbs_df.mean(axis=1)

# Display the results
print("Coefficient of Variation for vvDD condition:")
print(vvDD_cv)

print("\nCoefficient of Variation for PBS condition:")
print(pbs_cv)

# %% [markdown]
*Since the CoV is increasing as the mean decreases for the vvDD condition, while almost constant for the PBS condition, we decide to use a addtive noise model adjusted by the signal intensity (tumor volume).*

# %% [markdown]
# Creation of the measurements_scaled.tsv file

# %% [code]
# make the measurement.tsv file
def create_rows(df, condition):
    rows = []
    for time in df.index:
        for col in df.columns:
            rows.append({
                'observableId': 'tumor_num',
                'simulationConditionId': condition,
                'measurement': df.loc[time, col],
                'time': time,
            })
    return rows

# Create rows for vvDD and PBS conditions
vvDD_rows = create_rows(scaled_tumor_vol_vvDD_df, 'vvDD')
pbs_rows = create_rows(scaled_tumor_vol_pbs_df, 'ctrl')

# Combine all rows
all_rows = vvDD_rows + pbs_rows

# Create the final dataframe
measurements_df = pd.DataFrame(all_rows)

# Reorder columns to match the specified order
column_order = ['observableId', 'simulationConditionId', 'measurement', 'time']
measurements_df = measurements_df[column_order]

# Save to TSV file
measurements_df.to_csv('petab_files/measurements_scaled.tsv', sep='\t', index=False)

# %% [markdown]
# Creation of parameters.tsv, observable.tsv, conditions.tsv, visualization.tsv file

# %% [code]
# Create the parameter table
parameter_df = pd.DataFrame({
    'parameterId': ['rho', 'kappa', 'psi', 'phi', 'beta', 'alpha', 'delta', 'sigma_a', 'sigma_b'],
    'parameterName': ['rho', 'kappa', 'psi', 'phi', 'beta', 'alpha', 'delta', 'sigma_a', 'sigma_b'],
    'parameterScale': ['log10'] * 9,
    'lowerBound': [0.42, 1e2, 1e-10, 1e-8, 1e0, 1e-4, 1e-2, 1e1, 1e-4],
    'upperBound': [1.66, 1e5, 1e-2, 1e8, 1e4, 1e3, 1e2, 1e5, 1e2],
    'nominalValue': [1] * 9,
    'estimate': [1] * 9,
    'parameterType': [''] * 9
})
# export the parameter table to tsv file
parameter_df.to_csv('petab_files/parameters.tsv', sep='\t', index=False)


# %% [code]
# Create the observable table
L = 5

observable_formula = 'U + ' + ' + '.join([f'I_{j}' for j in range(1, L + 1)])
noise_formula = 'sqrt(sigma_a^2 + (sigma_b * (U + ' + ' + '.join([f'I_{j}' for j in range(1, L + 1)]) + '))^2)'

observable_df = pd.DataFrame({
    'observableId': ['tumor_num'],
    'observableName': ['tumor_number'],
    'observableFormula': [observable_formula],
    'noiseFormula': [noise_formula],
    'noiseDistribution': ['normal'],
    'observableTransformation': ['lin']
})

# export the observable table to tsv file
observable_df.to_csv('petab_files/observables.tsv', sep='\t', index=False)

# %% [code]
# Create the condition table
condition_df = pd.DataFrame({
    'conditionId': ['vvDD', 'ctrl'],
    'conditionName': ['vvDD', 'ctrl'],
    'u_2': [1e9, 0]
})
# export the condition table to tsv file
condition_df.to_csv('petab_files/conditions.tsv', sep='\t', index=False)

# %% [code]
# Create the visualization table

visualization_df = pd.DataFrame({
    'plotId': ['plot1'],
    'plotTypeData': ['MeanAndSD'],
    'plotTypeSimulation': ['ScatterPlot'],
    'xValues': ['condition'],
    'xLabel': ['Condition'],
    'yValues': ['tumor_num'],
    'yLabel': ['Tumor Number'],
    'legendEntry': ['Model']
})

# export the visualization table to tsv file
visualization_df.to_csv('petab_files/visualizations.tsv', sep='\t', index=False)

# %% [code]
```
