## Supplementary material for "Parameter Estimation and Model Selection for the Quantitative Analysis of Oncolytic Virus Therapy in Zebrafish^⋆^": All additional figures and results, together with all SBML and PEtab files, and the code and instructions to perform all analyses: README.md

## For age-of-infection model with scaling parameter:

```
------------------------ 
 >>> RESULTS SUMMARY:
 ------------------------ 

 Observability-Identifiability matrix calculated in 6.039259e+01 seconds. 

 >>> The rank of the observability matrix is 21 
     transcendence degree of k(U,Y) --> k(U,Y,x,p) is 1 

 >>> The model is not FISPO:
 >>> These parameters are identifiable:
      [rho_, psi_, phi_, alpha_] 
 >>> These parameters are unidentifiable:
      [kappa_, beta_, delta_, s_] 
 >>> These states are observable (and their initial conditions, if considered unknown, are identifiable):
      [VExp1, VExp2] 
 >>> These states are unobservable (and their initial conditions, if considered unknown, are unidentifiable):
      [UExp1, I_1Exp1, I_2Exp1, I_3Exp1, I_4Exp1, I_5Exp1, UExp2, I_1Exp2, I_2Exp2, I_3Exp2, I_4Exp2, I_5Exp2] 
 >>> Matrix analyzed in 6.563950e-02 seconds. 

 Total execution time: 6.045823e+01 seconds. 
 ```

## For age-of-infection model without scaling parameter:

 ```
------------------------ 
 >>> RESULTS SUMMARY:
 ------------------------ 

 Observability-Identifiability matrix calculated in 5.838756e+01 seconds. 

 >>> The rank of the observability matrix is 21 
     transcendence degree of k(U,Y) --> k(U,Y,x,p) is 0 

 >>> The model is FISPO.
     All its states are observable.
     All its parameters are locally structurally identifiable.
 >>> Matrix analyzed in 2.600208e-02 seconds. 

 Total execution time: 5.841357e+01 seconds. 
 ```
