## Supplementary material for "Parameter Estimation and Model Selection for the Quantitative Analysis of Oncolytic Virus Therapy in Zebrafish^⋆^": All additional figures and results, together with all SBML and PEtab files, and the code and instructions to perform all analyses: model_creation.py

```
import tellurium as te
import libsbml

# Model definition
baseline_model = """
model baseline_model

  // ODE functions
  U' = rho * U * (1 - (U + I)/kappa) - psi * V * U;
  I' = rho * I * (1 - (U + I)/kappa) + psi * V * U - alpha * I;
  V' = alpha * beta * I - psi * V * U - delta * V;

  // initial conditions
  // number of uninfected tumor cells at t=0, when the virus is injected
  U = kappa / (1 + (kappa/400 - 1) * exp(-rho));   // u_1 = 400, the number of tumor cells injected at t=-1
  // number of infected tumor cells at t=0, when the virus is injected
  I = 0; 
  // number of virus injected at t=0
  V = u_2; 

  // condition dependent parameters, 0 or 1e9, which can be obtained from the petab
  u_2 = 1 * 1E9; // pfu
  
  // parameters to be estimated
  rho = 0.01;
  kappa = 0.01;
  psi = 0.01;
  alpha = 0.01;
  beta = 0.01;
  delta = 0.01;

  // Event to set U to zero when U falls below 1e-8, to avoid exploding solutions that are not biologically relevant
  at (U < 1e-8): U = 0;

end
"""

# Load the model
r = te.loada(baseline_model)
# Export the model to SBML
r.exportToSBML('petab_files/baseline_model.xml', current=False)
```
