## Supplementary material for "Parameter Estimation and Model Selection for the Quantitative Analysis of Oncolytic Virus Therapy in Zebrafish^⋆^": All additional figures and results, together with all SBML and PEtab files, and the code and instructions to perform all analyses: README.md

## For baseline model with the sclaing parameter:

```
 ------------------------ 
 >>> RESULTS SUMMARY:
 ------------------------ 

 Observability-Identifiability matrix calculated in 7.921966e+00 seconds. 

 >>> The rank of the observability matrix is 12 
     transcendence degree of k(U,Y) --> k(U,Y,x,p) is 1 

 >>> The model is not FISPO:
 >>> These parameters are identifiable:
      [rho_, alpha_, beta_, delta_] 
 >>> These parameters are unidentifiable:
      [kappa_, psi_, s_] 
 >>> These states are unobservable (and their initial conditions, if considered unknown, are unidentifiable):
      [UExp1, IExp1, VExp1, UExp2, IExp2, VExp2] 
 >>> Matrix analyzed in 3.789821e-02 seconds. 

 Total execution time: 7.959864e+00 seconds. 
```

## For baseline model without the scaling parameter:

```
------------------------ 
 >>> RESULTS SUMMARY:
 ------------------------ 

 Observability-Identifiability matrix calculated in 7.692430e+00 seconds. 

 >>> The rank of the observability matrix is 12 
     transcendence degree of k(U,Y) --> k(U,Y,x,p) is 0 

 >>> The model is FISPO.
     All its states are observable.
     All its parameters are locally structurally identifiable.
 >>> Matrix analyzed in 2.035946e-02 seconds. 

 Total execution time: 7.712789e+00 seconds. 
```
