## Supplementary material for "Parameter Estimation and Model Selection for the Quantitative Analysis of Oncolytic Virus Therapy in Zebrafish^⋆^": All additional figures and results, together with all SBML and PEtab files, and the code and instructions to perform all analyses: visualization.ipynb

```
# %% [code]
import numpy as np
import pandas as pd
import scipy
import matplotlib.pyplot as plt
import matplotlib
import statsmodels.api as sm
import os

import petab
import pypesto
import pypesto.petab
from pypesto.optimize import minimize
from pypesto.engine import MultiProcessEngine
from pypesto.optimize.optimizer import FidesOptimizer
from pypesto.visualize import waterfall, parameters, profiles
from pypesto.visualize.model_fit import _get_simulation_rdatas
import pypesto.profile as profile
from pypesto.optimize import ScipyOptimizer
from pypesto.profile.options import ProfileOptions

# %% [markdown]
Some plotting helper functions

# %% [code]
def hex_to_rgba_gradient(color1, color2, n):
    '''
    Create a gradient in rgba between two hex colors
    '''
    # Convert to rgba
    c1 = matplotlib.colors.to_rgba(matplotlib.colors.hex2color(color1))
    c2 = matplotlib.colors.to_rgba(matplotlib.colors.hex2color(color2))

    return [[(c1[i]*(n-j-1) + c2[i]*j)/(n-1) for i in range(4)] for j in range(n)]

# find the index for cut off based on Chi square distribution CI 95%
def find_cut_off_index(result, ci = 0.95):
    '''
    Find the cut off index for the data based on the Chi square distribution
    '''

    # calculate the chi square distribution
    cut_off_value = scipy.stats.chi2.ppf(ci, 1)

    # find the index
    best_fval = result.optimize_result.list[0].fval

    for i in range(len(result.optimize_result.list)):
        if result.optimize_result.list[i].fval > best_fval + cut_off_value:
            break
    
    return i - 1

# %% [code]
# Plot setting
plt.rcParams['font.size'] = 30
plt.rcParams['font.family'] = 'Arial'

dpi = 100
wid = int(2560/dpi)
hei = int(1600/dpi)

red_color = '#f78884'
blue_color = '#06688c'

# Define the folder where you want to save the figures
folder_path = "../../figure/individual_based_age_of_infection_model/"
# If the folder does not exist, create it
if not os.path.exists(folder_path):
    os.makedirs(folder_path)

# %% [markdown]
# import the petab problem and load the optimization result

# %% [code]
# number of optimization runs
n_runs, max_iter = 5000, 5000
# set the random seed
np.random.seed(500)

# %% [code]
petab_yaml = 'petab_files/individual_based_age_of_infection_model.yaml'
petab.validate(petab_yaml)
petab_problem = petab.Problem.from_yaml(petab_yaml)

problem = pypesto.petab.PetabImporter(
        petab_problem,
        hierarchical=False,
        model_name=f"Individual_Based_age_of_Infection_Model",
    ).create_problem(force_compile=True, verbose=False)
problem.objective.amici_model.setAllStatesNonNegative()

# some model properties
print("Model parameters:", list(problem.objective.amici_model.getParameterIds()), "\n")
print("Model const parameters:", list(problem.objective.amici_model.getFixedParameterIds()), "\n")
print("Model outputs:   ", list(problem.objective.amici_model.getObservableIds()), "\n")
print("Model states:    ", list(problem.objective.amici_model.getStateIds()), "\n")

# %% [code]
# load result history from file
result = pypesto.store.read_result('optimization_history/individual_based_age_of_infection_model.hdf5')

# print result summary
print(result.summary())

# %% [code]
parameters_from_result = dict(zip(problem.x_names, result.optimize_result.list[0]['x']))
# Scale all parameters and put them into a dictionary
scaled_parameters = {key: 10**value for key, value in parameters_from_result.items()}

# Print the scaled parameters
print("Scaled parameters:")
for key, value in scaled_parameters.items():
    print(f"{key}: {value}")

# %% [markdown]
# Obtain data and visualize the fitting result

# %% [code]
return_dict = problem.objective(result.optimize_result.list[0].x, return_dict=True)
rdatas = return_dict['rdatas']
edatas = problem.objective.edatas
x_axis = [edata.id for edata in edatas]
simulation = [rdata.y.reshape(5, -1)[:,0] for rdata in rdatas]
data = [np.array(edata.getObservedData()) for edata in edatas]

# %% [code]
# get the statistics for 95% CI
cut_off_index = find_cut_off_index(result)
# define as in petab_files_creation.ipynb to convert the data back to tumor volume
s = 3510.7678534742176

# %% [code]
"""
visualize the temporal dynamics of the virus, uninfected and infected tumor cells using the fitted model from the result
from day 3 to day 7
get the simulation results for the optimized parameters
"""

amici_model = problem.objective.amici_model

L = 5
species_to_plot = ['U', 'I_1', f'I_{L}', 'V_e']

# simulate from day 3 to day 12
stop_day = 4
timepoints = np.linspace(start=0, stop=stop_day, num=50)

simulation_rdatas = _get_simulation_rdatas(
    result=result,
    problem=problem,
    start_index = 0,
    simulation_timepoints=timepoints,
)


# %% [code]
# Create a figure with subplots
fig, axs = plt.subplots(10, 2, figsize=(8, 25))

# for calculation of 1-sigma for the simulation
sigma_a = scaled_parameters['sigma_a']
sigma_b = scaled_parameters['sigma_b']

# Plot the data and simulation for each individual
for i in range(20):
     row = i % 10
     col = 0 if i < 10 else 1
     color = red_color if i < 10 else blue_color

     axs[row, col].plot(np.array([3,4,5,6,7]), data[i]*s, 
                        marker='o', markersize=8, lw=5, linestyle='--', color=color, alpha=0.5, label='control (data)' if i < 10 else 'vvDD (data)')
     axs[row, col].plot(timepoints+3, simulation_rdatas[i]['y']*s,
                            linestyle='-', lw=5, color=color, alpha=1, label='control (simulation)' if i < 10 else 'vvDD (simulation)')
    
     sigma_ = np.sqrt(sigma_a**2 + (simulation_rdatas[i]['y'].reshape(1, -1)[0] * sigma_b)**2)
     axs[row, col].fill_between(timepoints+3, (simulation_rdatas[i]['y'].reshape(1, -1)[0] - sigma_)*s, (simulation_rdatas[i]['y'].reshape(1, -1)[0] + sigma_)*s, 
             color=color, alpha=0.2, edgecolor='none')

     axs[row, col].set_title(rf'$D_c^{{{i+1}}}$' if i < 10 else rf'$D_v^{{{i-9}}}$', fontsize=30, color=color)
     axs[row, col].grid(False)
     axs[row, col].set_xticks([3, 4, 5, 6, 7])
     axs[row, col].spines['top'].set_visible(False)
     axs[row, col].spines['right'].set_visible(False)
     axs[row, col].spines['bottom'].set_linewidth(2)
     axs[row, col].spines['left'].set_linewidth(2)
     # Make y-axis labels more sparse
     axs[row, col].yaxis.set_major_locator(plt.MaxNLocator(2))
     # Make ticks thicker
     axs[row, col].tick_params(width=2)

# Adjust the position of the y-axis labels
fig.text(0.04, 0.5, r'Tumor Volume [$\mu m^3$]', va='center', rotation='vertical', fontsize=30)
fig.text(0.52, 0.04, 'Time [days]', ha='center', fontsize=30)

# Add legends
handles1, labels1 = axs[0, 0].get_legend_handles_labels()
handles3, labels3 = axs[0, 1].get_legend_handles_labels()

# Separate red and blue labels
red_handles = [handles1[1], handles1[0]]
red_labels = [labels1[1], labels1[0]]
blue_handles = [handles3[1], handles3[0]]
blue_labels = [labels3[1], labels3[0]]

fig.legend(red_handles + blue_handles, red_labels + blue_labels, loc='lower center', ncol=2, frameon=False, fontsize=20, bbox_to_anchor=(0.6, -0.025))

plt.tight_layout(rect=[0.05, 0.05, 1, 0.95])
plt.savefig(folder_path + 'individual_trajectory_complete.pdf', dpi=300, bbox_inches='tight')
plt.show()


# %% [code]
fig = plt.figure(figsize=(8, 6))

gs = fig.add_gridspec(2, 2)
ax1 = fig.add_subplot(gs[0, 0])
ax2 = fig.add_subplot(gs[1, 0])
ax3 = fig.add_subplot(gs[0, 1])
ax4 = fig.add_subplot(gs[1, 1])

# calculate the 1-sigma for the simulation
sigma_a = scaled_parameters['sigma_a']
sigma_b = scaled_parameters['sigma_b']

# Panel b: State trajectories
axs = [ax1, ax2, ax3, ax4]
for j, i in enumerate([6, 7, 10, 11]):
    condition = 'ctrl'  if i < 10 else 'vvDD'
    ax = axs[j]
    color = red_color if condition == 'ctrl' else blue_color
    ax.plot(np.array([3,4,5,6,7]), data[i]*s, 
                        marker='o', markersize=8, lw=5, linestyle='--', color=red_color if i < 10 else blue_color, alpha=0.5, label='control (data)' if i < 10 else 'vvDD (data)')
    ax.plot(timepoints+3, simulation_rdatas[i]['y']*s,
                            linestyle='-', lw=5, color=red_color if i < 10 else blue_color, alpha=1, label='control (simulation)' if i < 10 else 'vvDD (simulation)')
    
    sigma_ = np.sqrt(sigma_a**2 + (simulation_rdatas[i]['y'].reshape(1, -1)[0] * sigma_b)**2)
    ax.fill_between(timepoints+3, (simulation_rdatas[i]['y'].reshape(1, -1)[0] - sigma_)*s, (simulation_rdatas[i]['y'].reshape(1, -1)[0] + sigma_)*s, 
             color=color, alpha=0.2, edgecolor='none')
    
    ax.set_xticks(np.arange(3, stop_day + 4, 1))
    ax.set_xticklabels(np.arange(3, stop_day + 4, 1))
    if i == 6:
        ax.set_title(r'$D_c^7$', fontsize=30, color=red_color)
    if i == 7:
        ax.set_title(r'$D_c^8$', fontsize=30, color=red_color)
    if i == 10:
        ax.set_title(r'$D_v^1$', fontsize=30, color=blue_color)
    if i == 11:
        ax.set_title(r'$D_v^2$', fontsize=30, color=blue_color)
    ax.yaxis.set_tick_params(labelleft=True)
    ax.spines['top'].set_visible(False)
    ax.spines['right'].set_visible(False)
    ax.spines['left'].set_linewidth(2)
    ax.spines['bottom'].set_linewidth(2)
    ax.tick_params(width=2)

# Adjust the position of the y-axis labels
fig.text(-0.06, 0.5, r'Tumor Volume [$\mu m^3$]', va='center', rotation='vertical', fontsize=30)
fig.text(0.5, -0.04, 'Time [days]', ha='center', fontsize=30)

# ax1.text(-0.2, 1.3, 'a', transform=ax1.transAxes, fontsize=40, fontweight='bold', va='top', ha='right')
# ax3.text(-0.05, 1.3, 'b', transform=ax3.transAxes, fontsize=40, fontweight='bold', va='top', ha='right')

handles1, labels1 = ax1.get_legend_handles_labels()
handles3, labels3 = ax4.get_legend_handles_labels()

# Separate red and blue labels
red_handles = [handles1[1], handles3[1]]
red_labels = [labels1[1], labels3[1]]
blue_handles = [handles1[0], handles3[0]]
blue_labels = [labels1[0], labels3[0]]

fig.legend(red_handles + blue_handles, red_labels + blue_labels, loc='center right', ncol=1, frameon=False, fontsize=30, bbox_to_anchor=(1.65, 0.5))

plt.subplots_adjust(wspace=0.7, hspace=0.1)  # Adjust space between subplots

plt.tight_layout()
plt.savefig(folder_path + 'individual_trajectory.png', dpi=300, bbox_inches='tight')
plt.show()

# %% [markdown]
# Parameter estimation analysis

# %% [code]
result.problem.x_scales = ['log10'] * len(result.problem.x_names)

# %% [code]
panel_labels = ['a', 'b', 'c', 'd']

# %% [code]
plt.rcParams.update({'font.size': 30})

fig = plt.figure(figsize=(wid*1.15, hei*1.75))

gs = fig.add_gridspec(5, 3)
ax1 = fig.add_subplot(gs[0:2, :])
ax2 = fig.add_subplot(gs[2:, 0])
ax3 = fig.add_subplot(gs[2:, 1])
ax4 = fig.add_subplot(gs[2:, 2])

waterfall(result, ax=ax1)
ax1.set_ylabel('Objective value')

# Parameter plots in the second row
pypesto.visualize.parameters(result, ax=ax2, plot_inner_parameters=False, start_indices=cut_off_index, colors=hex_to_rgba_gradient('#A7C9F8', '#28518B', cut_off_index))
pypesto.visualize.parameters(result, ax=ax3, plot_inner_parameters=False, start_indices=300, colors=hex_to_rgba_gradient('#A7C9F8', '#28518B', 300))
pypesto.visualize.parameters(result, ax=ax4, plot_inner_parameters=False, start_indices=100, colors=hex_to_rgba_gradient('#A7C9F8', '#28518B', 100))

ax2.set_title('95% CI', fontsize=30)
ax3.set_title('Top 300', fontsize=30)
ax4.set_title('Top 100', fontsize=30)

# Set all the x-axis, x and y labels to have fontsize 30
for ax in [ax2, ax3, ax4]:
    ax.set_xticklabels(ax.get_xticklabels(), fontsize=20)
    ax.set_xlabel('Parameter Value', fontsize=30)
ax2.set_ylabel('Parameter', fontsize=30)

# Remove top and right lines and make lines and ticks thicker
for i, ax in enumerate([ax1, ax2, ax3, ax4]):
    ax.spines['top'].set_visible(False)
    ax.spines['right'].set_visible(False)
    ax.spines['left'].set_linewidth(2)
    ax.spines['bottom'].set_linewidth(2)
    ax.tick_params(width=2)
    # Add panel labels
    if i == 0:
        ax.text(-0.115, 1.05, panel_labels[i], transform=ax.transAxes, fontsize=40, fontweight='bold', va='top', ha='right')
    else:
        ax.text(-0.55, 1.05, panel_labels[i], transform=ax.transAxes, fontsize=40, fontweight='bold', va='top', ha='right')

plt.tight_layout()
plt.savefig(os.path.join(folder_path, 'waterfall_parameters_plot.pdf'), dpi=dpi, bbox_inches="tight")
plt.show()

# %% [markdown]
# Profiling

# %% [code]
result = profile.parameter_profile(
    problem=problem,
    result=result,
    engine=MultiProcessEngine(),
    optimizer=ScipyOptimizer(),
    result_index=0,
    profile_options=ProfileOptions(whole_path=True),
)

# %% [code]
from pypesto.profile import calculate_approximate_ci, chi2_quantile_to_ratio

# extract problem
problem = result.problem
# extract profile list
profile_list = result.profile_result.list[0]

confidence_ratio = chi2_quantile_to_ratio(0.95)

# calculate intervals
intervals = []
for i_par in range(problem.dim_full):
    xs = profile_list[i_par].x_path[i_par]
    ratios = profile_list[i_par].ratio_path
    lb, ub = calculate_approximate_ci(
        xs=xs, ratios=ratios, confidence_ratio=confidence_ratio
    )
    intervals.append((lb, ub))

# %% [code]
fig = plt.figure(figsize=(wid*1.15, hei*1.75))

labels = ["$\\rho$", "$\\kappa$", "$\\psi$", "$\\phi$", "$\\beta$", "$\\alpha$", "$\\delta$"] + [f"$\\xi_{{{i}}}$" for i in range(1,21)] + ["$\\sigma_{a}$", "$\\sigma_{b}$"]

ax = profiles(result, show_bounds=True, size=(50, 30), colors=[0,0,0,1], profile_list_ids=len(result.profile_result.list)-1, quality_colors=False)

cut_off_value = scipy.stats.chi2.ppf(0.95, 1)  # 95% confidence interval cut-off value

for i, a in enumerate(ax):
    a.set_ylim([0, 1.1])
    a.spines['top'].set_visible(False)
    a.spines['right'].set_visible(False)
    a.spines['left'].set_linewidth(2)
    a.spines['bottom'].set_linewidth(2)
    a.tick_params(width=2, labelsize=25)  # Set tick label size to 20
    for label in a.get_xticklabels():
        label.set_rotation(45)
    # a.set_xlabel(rf'{labels[i]}', fontsize=40)  # Change x-axis label to math notation and increase font size
    a.set_xlabel('')  # Remove innate x-label
    a.set_ylabel('')  # Remove innate y-label
    if len(a.get_xticks()) == 12:
        a.set_xticks(a.get_xticks()[::3])
    if len(a.get_xticks()) == 9 or len(a.get_xticks()) == 8 or len(a.get_xticks()) == 7:
        a.set_xticks(a.get_xticks()[::2])
    if i == 0:
        a.set_xticklabels([f'${{{10**(tick):.2f}}}$' for tick in a.get_xticks()])
    else:
        a.set_xticklabels([f'$10^{{{int(tick)}}}$' for tick in a.get_xticks()])
    # Add red dashed line for cut-off threshold
    a.axhline(y=np.exp(-cut_off_value/2), color='blue', linestyle='--', linewidth=2)
    # Add vertical lines for confidence interval bounds
    lb, ub = intervals[i]
    # Add horizontal line for confidence interval bounds
    a.hlines(y=np.exp(-cut_off_value/2), xmin=lb, xmax=ub, color='red', linestyle='-', linewidth=2)
    # Add small vertical bars at the bounds if they are not at the boundary of the axis
    if lb > a.get_xlim()[0]:
        a.vlines(x=lb, ymin=np.exp(-cut_off_value/2) - 0.02, ymax=np.exp(-cut_off_value/2) + 0.02, color='red', linestyle='-', linewidth=3)
    if ub < a.get_xlim()[1]:
        a.vlines(x=ub, ymin=np.exp(-cut_off_value/2) - 0.02, ymax=np.exp(-cut_off_value/2) + 0.02, color='red', linestyle='-', linewidth=3)
    # Add annotation of the letter "x" to the bottom right of each subfigure
    a.annotate(rf'{labels[i]}', xy=(0.2, 0.25), xycoords='axes fraction', fontsize=40, ha='right', va='bottom')

# Add a common y-label
fig.text(0.03, 0.5, 'Likelihood Ratio', va='center', rotation='vertical', fontsize=30)
# Adjust the position of the y-axis labels
for a in ax:
    a.xaxis.set_label_coords(0.55, -0.5)

plt.subplots_adjust(wspace=0.3, hspace=0.5)
plt.savefig(os.path.join(folder_path, 'profile_plot_res.pdf'), dpi=dpi, bbox_inches="tight")
plt.show()

# %% [markdown]
# AIC and QQ plot

# %% [code]
# Calculate AIC
AIC = 2 * len(result.optimize_result.list[0]['x']) + 2 * result.optimize_result.list[0]['fval']

print(f"AIC: {AIC}")

# %% [code]
n_measurements = 100
n_parameters = len(result.optimize_result.list[0]['x'])

# Calculate AIC for the small dataset
AIC_small = AIC + (2. * n_parameters**2 + 2.*n_parameters)/(n_measurements - n_parameters - 1)

print(f"AICc: {AIC_small}")

# %% [code]
import statsmodels.api as sm
import seaborn as sns

from statsmodels.stats.stattools import durbin_watson
from statsmodels.stats.diagnostic import het_breuschpagan

# %% [code]
# Your data arrays
empirical_data = np.array(data)
simulation_data = np.array(simulation)

# Calculate residuals
residuals = empirical_data - simulation_data

# Residue plot
for i in range(5):
    fig, ax = plt.subplots(figsize=(10,8))
    ax.scatter(np.arange(20), residuals[:,i], alpha=1)
    ax.axhline(y=0, color='r', lw=3, linestyle='--')
    ax.set_xlabel('Index')
    ax.set_ylabel('Residuals')
    ax.set_title('Residual Plot')
    ax.spines['top'].set_visible(False)
    ax.spines['right'].set_visible(False)
    ax.spines['bottom'].set_linewidth(2)
    ax.spines['left'].set_linewidth(2)
    ax.grid(False)
    plt.tight_layout()
    # plt.savefig(folder_path + 'residual_plot.pdf', dpi=300, bbox_inches='tight')
    plt.show()

# %% [code]
# Create QQ plot
fig, ax = plt.subplots(figsize=(wid, hei))
sm.qqplot_2samples(sm.ProbPlot(empirical_data), sm.ProbPlot(simulation_data), line='45', ax=ax)
ax.set_xlabel('Empirical Data Quantiles')
ax.set_ylabel('Simulation Data Quantiles')
ax.set_title('QQ Plot: Empirical Data vs. Simulation Results')
ax.grid(False)
ax.spines['top'].set_visible(False)
ax.spines['right'].set_visible(False)
ax.spines['bottom'].set_linewidth(2)
ax.spines['left'].set_linewidth(2)
plt.tight_layout()
# plt.savefig(folder_path + 'qq_plot.pdf', dpi=300, bbox_inches='tight')
plt.show()


# %% [code]
```
