## Supplementary material for "Parameter Estimation and Model Selection for the Quantitative Analysis of Oncolytic Virus Therapy in Zebrafish^⋆^": All additional figures and results, together with all SBML and PEtab files, and the code and instructions to perform all analyses: Lipschitz_Continuity_Proof.pdf

### Proof of Lipschitz Continuity for All Models*

**3 Individual-based age-of-infection model** 4

### 1 Baseline model

Here we show that the **baseline model** shown below is Lipschitz continuous with respect to the state variables on any bounded domain in $\mathbb{R}^3$.

#### 1.1 Model definition

Let $X = (U, I, V) \in \mathbb{R}^3$. Define the vector field, $F(X) = (F_U, F_I, F_V)$, with

$$\begin{aligned} F_U(U, I, V) &= \rho U \left(1 - \frac{U+I}{\kappa}\right) - \psi V U, \\ F_I(U, I, V) &= \rho I \left(1 - \frac{U+I}{\kappa}\right) + \psi V U - \alpha I, \\ F_V(U, I, V) &= \alpha\beta I - \psi V U - \delta V. \end{aligned}$$

where $\rho, \psi, \alpha, \beta, \delta, \kappa$ are constants.

#### 1.2 Proof

##### 1.2.1 Examine the Functional Forms

Each right-hand side is composed of terms that are at most bilinear or quadratic in the variables of $X$. For example:

- $\rho U(1 - \frac{U+I}{\kappa})$ expands to $\rho U - \frac{\rho}{\kappa} U(U + I)$, which is a polynomial (up to quadratic) in $U$ and $I$.

*Supplementary of the paper: *Parameter Estimation and Model Selection for the Quantitative Analysis of Oncolytic Virus Therapy in Zebrafish*

- Terms like $\psi VU$ are bilinear in $V$ and $U$.
- Terms like $-\alpha I$ and $-\delta V$ are linear in their respective variables.

Importantly, there are no divisions by state variables (only by $\kappa$, which is constant) and no transcendental functions involved. Thus, each component $F_U$, $F_I$, and $F_V$ is a polynomial expression in $X$.

###### 1.2.2 Continuous differentiability

Since polynomials are infinitely differentiable, each component of $F$ is continuously differentiable with respect to $X$. Continuous differentiability implies local Lipschitz continuity.

###### 1.2.3 Bounded domain consideration

To show Lipschitz continuity on a specific set, consider any bounded domain $D \subset \mathbb{R}^3$. On $D$, there exists a constant $M > 0$ such that $\forall |x_i| \in X$,

$$x_i \leq M.$$

Because the partial derivatives of $F$ with respect to $X$ are at most linear in these variables, within a bounded domain they remain bounded. For example, the partial derivative $\frac{\partial F_U}{\partial U}$ looks like:

$$\frac{\partial F_U}{\partial U} = \rho - \frac{2\rho U}{\kappa} - \frac{\rho I}{\kappa} - \psi V.$$

Each of these is a polynomial in $X$ and thus is bounded on $D$.

###### 1.2.4 Existence of a lipschitz constant

Since all partial derivatives are bounded on the domain $D$, there exists a constant $L > 0$ such that $\forall X, Y \in D$:

$$\|F(X) - F(Y)\| \leq L\|X - Y\|.$$

This inequality satisfies the definition of Lipschitz continuity.

##### 1.3 Conclusion

The given system:

$$\begin{aligned}
\frac{dU}{dt} &= \rho U \left(1 - \frac{U + I}{\kappa}\right) - \psi VU, \\
\frac{dI}{dt} &= \rho I \left(1 - \frac{U + I}{\kappa}\right) + \psi VU - \alpha I, \\
\frac{dV}{dt} &= \alpha\beta I - \psi VU - \delta V
\end{aligned}$$

is continuously differentiable in $X = (U, I, V)$ and, therefore, Lipschitz continuous on any bounded domain in $\mathbb{R}^3$.

#### 2 Age-of-infection model

Here we show that the **age-of-infection model** shown below is Lipschitz continuous with respect to the state variables on any bounded domain in $\mathbb{R}^{L+2}$.

#### 2.1 Model definition

We denote the vector of state variables as $X = (U, I_1, I_2, \ldots, I_L, V) \in \mathbb{R}^{L+2}$ and $C = U + \sum_{l=1}^{L} I_l$.

Define the vector field $F(X) = (F_U, F_{I_1}, F_{I_2}, \ldots, F_{I_L}, F_V)$, where

$$\begin{aligned}
F_U &= \rho U\left(1 - \frac{C}{\kappa}\right) - \psi V U,\\
F_{I_1} &= \rho I_1\left(1 - \frac{C}{\kappa}\right) + \psi V U - \phi I_1,\\
F_{I_l} &= \rho I_l\left(1 - \frac{C}{\kappa}\right) + \phi(I_{l-1} - I_l) \quad \text{for } l \in \{2, \ldots, L-1\},\\
F_{I_L} &= \rho I_L\left(1 - \frac{C}{\kappa}\right) + \phi I_{L-1} - \alpha I_L,\\
F_V &= \beta\alpha I_L - \psi V U - \delta V.
\end{aligned}$$

#### 2.2 Proof

##### 2.2.1 Form of the functions

Each component of $F$ is composed of terms that are at most bilinear or affine of the variables in $X$. Specifically:

- $C = U + \sum_{l=1}^{L} I_l$ is a linear function of the state variables.
- The logistic-like terms, e.g., $\rho U(1 - \frac{C}{\kappa})$, expand to $\rho U - \frac{\rho}{\kappa} U(U + \sum I_l)$, which is a polynomial (quadratic) expression in the state variables.
- Terms like $\psi V U$, $\phi(I_{l-1} - I_l)$, and $\beta\alpha I_L$ are linear or bilinear in the state variables.

Since all these expressions are sums and products of the state variables and constants, each $F_i(X)$ is a polynomial (or at worst a polynomial-like function involving linear and bilinear terms) of the state variables.

##### 2.2.2 Continuous differentiability

Polynomials are infinitely differentiable functions. Hence, each component $F_i(X)$ is continuously differentiable with respect to all variables in $X$. Because $F$ is continuously differentiable, it follows that $F$ is locally Lipschitz continuous. This is a standard result in analysis: continuous differentiability implies local Lipschitz continuity.

##### 2.2.3 Boundedness of derivatives on a bounded domain

To be Lipschitz continuous on a specific domain, we must show that there exists a global Lipschitz constant on that domain. Consider any bounded subset $D \subset \mathbb{R}^{L+2}$. Since $D$ is bounded, there exists some $M > 0$ such that for all $X = (U, I_1, \ldots, I_L, V) \in D$, we have:

$$|U|, |I_l|, |V| \leq M \quad \text{for all } l = 1, \ldots, L.$$

Examine the Jacobian matrix $J_F(X)$ of $F$, whose entries are the partial derivatives $\frac{\partial F_i}{\partial x_j}$ where $x_j \in X$. Each partial derivative is again a linear or polynomial function of X. Because all variables are bounded by $M$ on $D$, each partial derivative is bounded by some constant that depends on $M$ and the parameters $\rho, \phi, \psi, \alpha, \beta, \delta, \kappa$.

Thus, there exists a constant $L > 0$ such that,

$$\forall X \in D, \|J_F(X)\| \leq L.$$

By the mean value theorem for vector-valued functions, if the Jacobian is bounded by $L$, then:

$$\|F(X) - F(Y)\| \leq L\|X - Y\| \quad \text{for all } X, Y \in D.$$

This shows that $F$ is Lipschitz continuous on the bounded domain $D$.

##### 2.3 Conclusion

We have demonstrated that:

- Each component of $F$ is continuously differentiable and thus locally Lipschitz.
- On any bounded domain, the partial derivatives of $F$ are bounded.

Therefore, $F$ is Lipschitz continuous on any bounded subset of $\mathbb{R}^{L+2}$.

#### 3 Individual-based age-of-infection model

It is trivial to show that the **individual-based age-of-infection model** is Lipschitz continuous by extending the age-of-infection model's initial conditions definition, thus not shown here.
