## Supplementary material for "Parameter Estimation and Model Selection for the Quantitative Analysis of Oncolytic Virus Therapy in Zebrafish^⋆^": All additional figures and results, together with all SBML and PEtab files, and the code and instructions to perform all analyses: README.md

# Parameter Estimation and Model Selection for the Quantitative Analysis of Oncolytic Virus Therapy in Zebrafish Embryos

<p align="center">
  <img src="figure/illustration/vvDD_mechanism.png" alt="Oncolytic Virus (vvDD) Mechanism" width="300">
</p>

## Overview
This repository contains the code and supplementary accompanying the paper, *Parameter Estimation and Model Selection for the Quantitative Analysis of Oncolytic Virus Therapy in Zebrafish Embryos (submitted to IFAC DYCOPS 2025)*

## Repository Contents
- **data/**: contains the original tumor volume measurements from [Mealiea et al. (2021)](https://www.nature.com/articles/s41417-020-0194-7.pdf)
- **figure/**: contains figure outputs from `visualization.ipynb` for each model, each subfolder containing:
    - model fits
    - profile plots
    - waterfall plot combined with parameter plots
- **model/**: contains the proof of Lipschitz continuity for all models and the following three models listed in the paper:
    - baseline model
    - age-of-infection model
    - individual-based age-of-infection model

in each model folder, there are files:

- `README.md`: structural identifiability information
- `model_creation.py`: create the `.xml` model file
- `petab_files_creation.ipynb`: build petab files defining the optimization problem
- `model_optimization.py`: perform optimization 
- `visualization.ipynb`: visualize the optimization results
- `check_gradients.ipynb`: double check the gradients of the model

## Requirement

After installing all the code using `git clone https://github.com/EchoRLiu/OV.git`, run the following command in the project folder to creat a virtual environment and install all the necessary packages

```
python -m venv env

source env/bin/activate

pip install -v -r requirements.txt

```
## Tutorial

**Quickstart**

To try one of the models,

1) first go to the model folder, e.g. individual_based_age_of_infection_model, and perform model optimization and save the printouts (if there is no `optimization_history` folder, please create one first to store all the results):

```
cd model/individual_based_age_of_infection_model

python model_optimization.py > optimization_history/result.out 2>&1

```

2) run through `visualization.ipynb` to visualize the results

3) (optional) run through the `check_gradients.ipynb` to check the gradients

## Contact
If you have any questions, please feel free to contact any of the authors:
- Yuhong Liu
- Dilan Pathirana
- Jan Hasenauer

or create an issue
