## Supplementary figures and images for "Parameter Estimation and Model Selection for the Quantitative Analysis of Oncolytic Virus Therapy in Zebrafish^⋆^"

### individual_trajectory.pdf

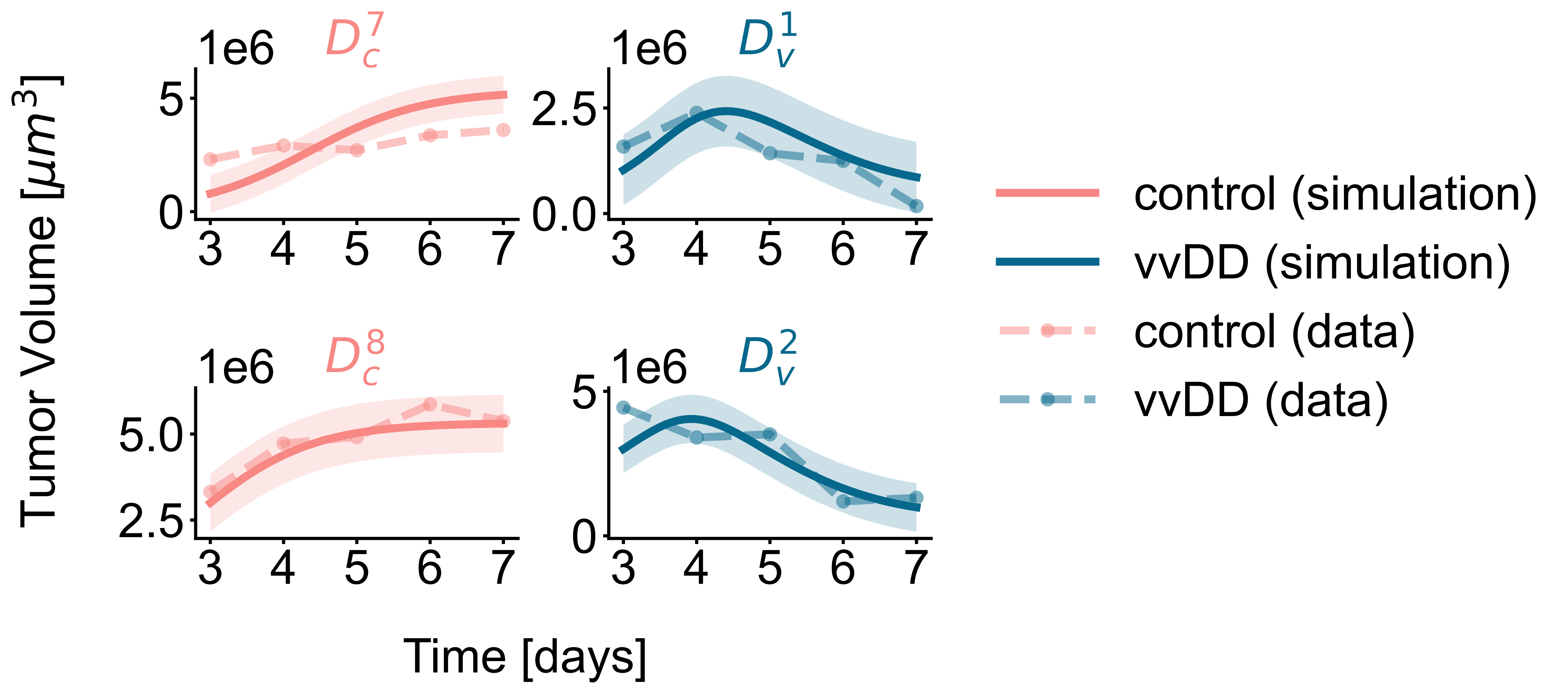

### individual_trajectory.png

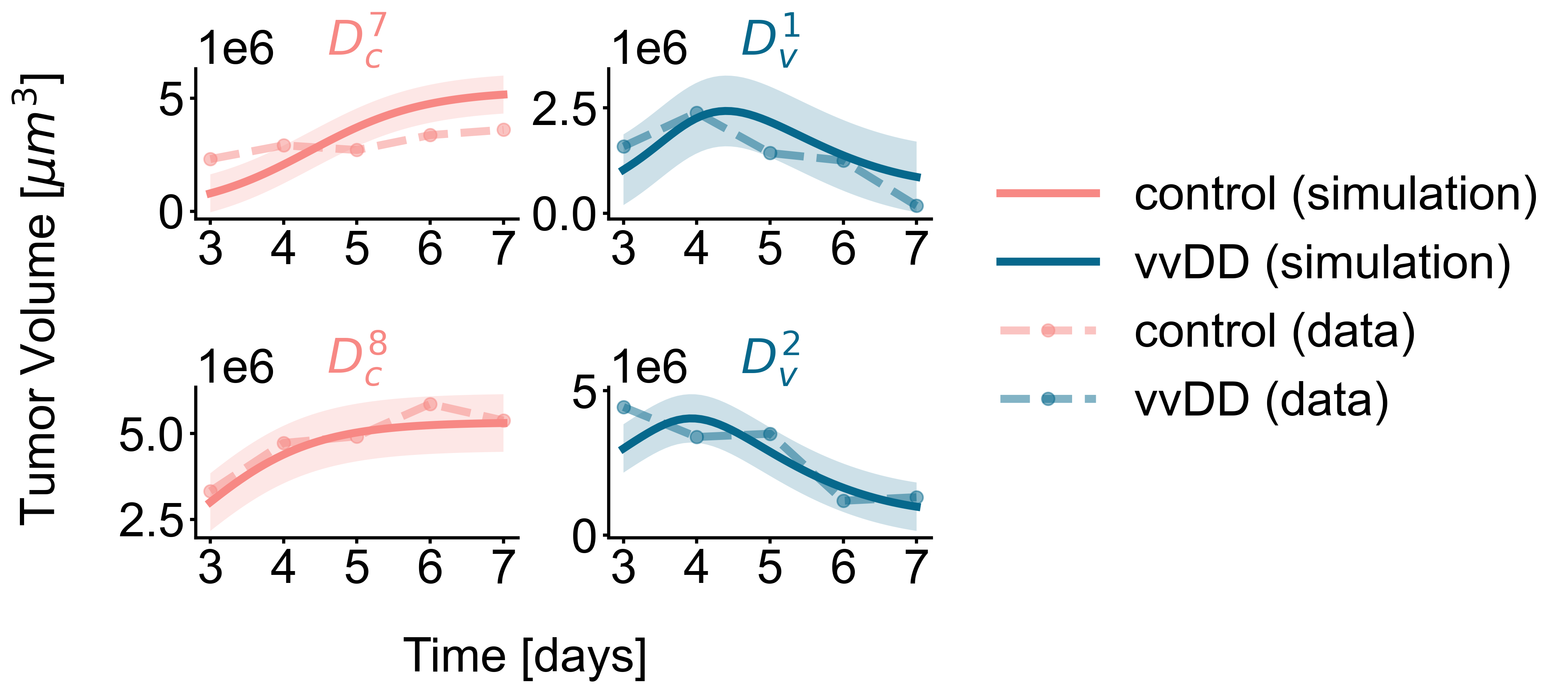

### individual_trajectory_complete.pdf

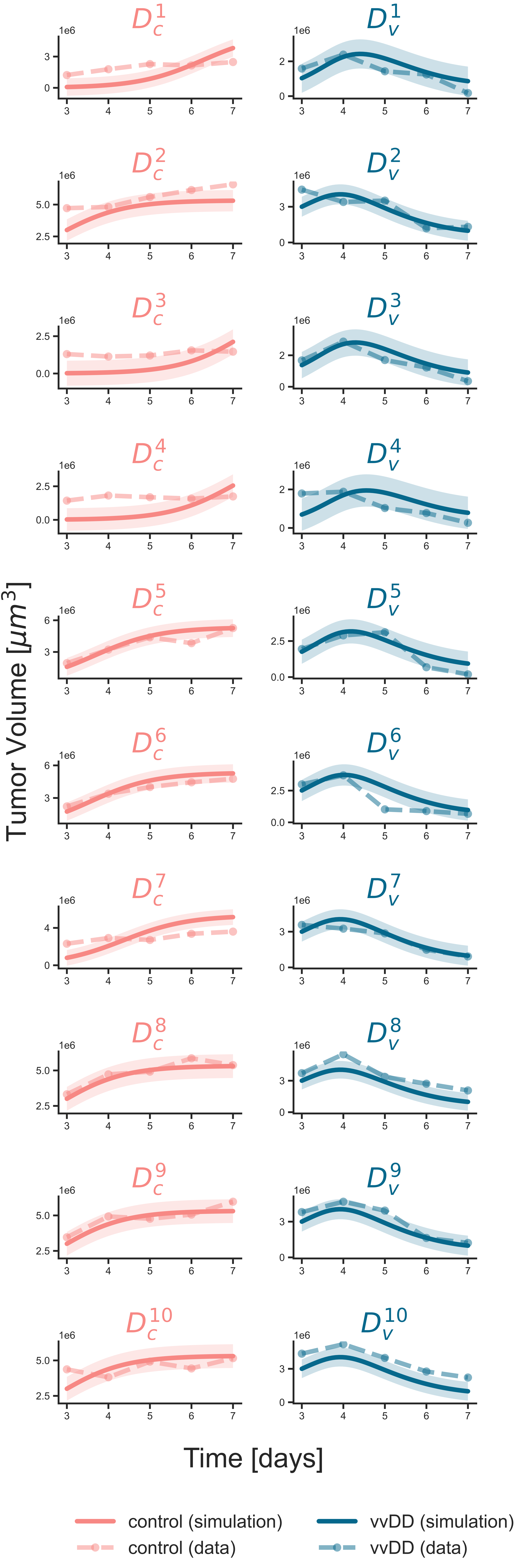

### original_data.pdf

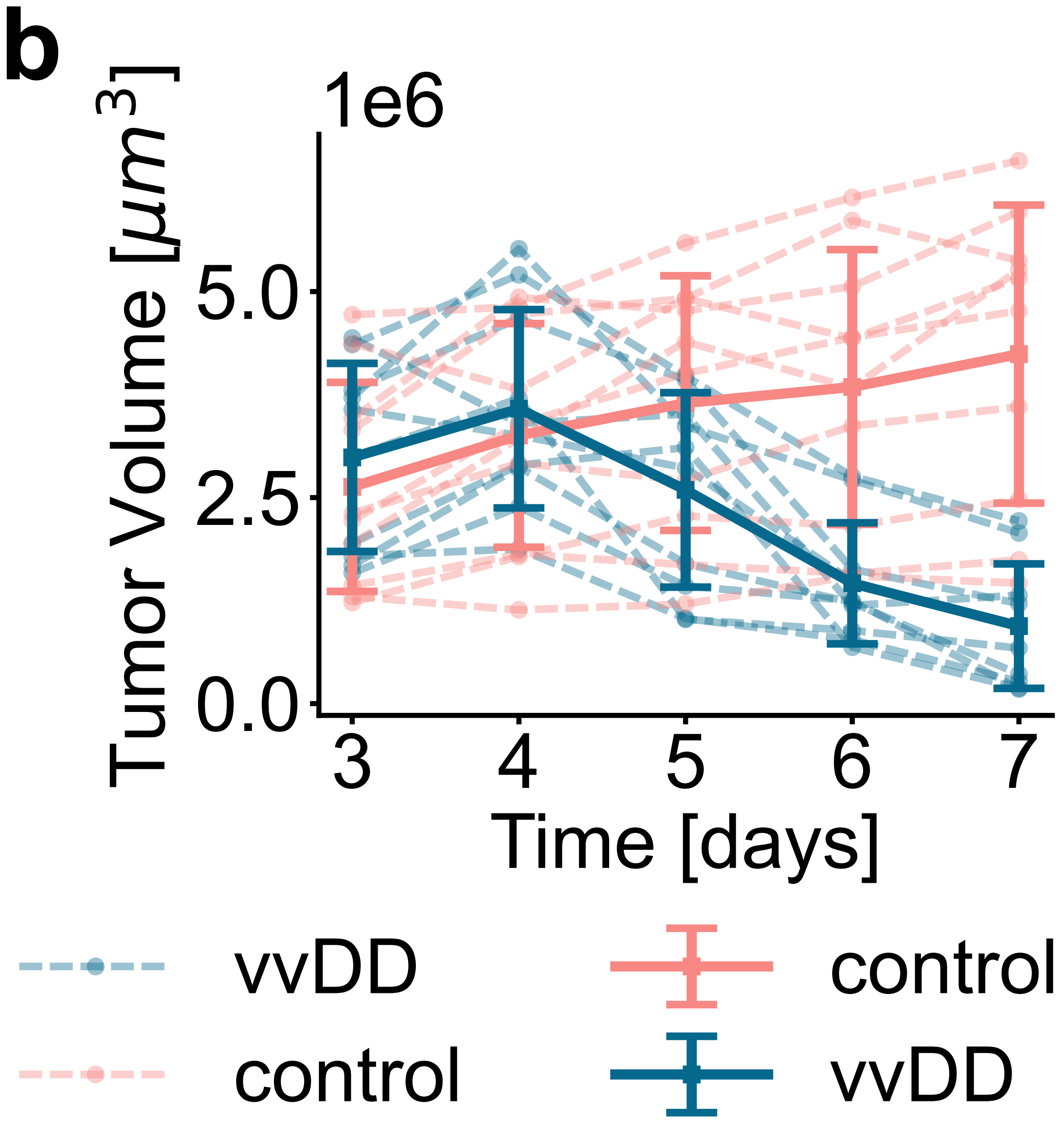

### original_imaging_and_quantification.jpg

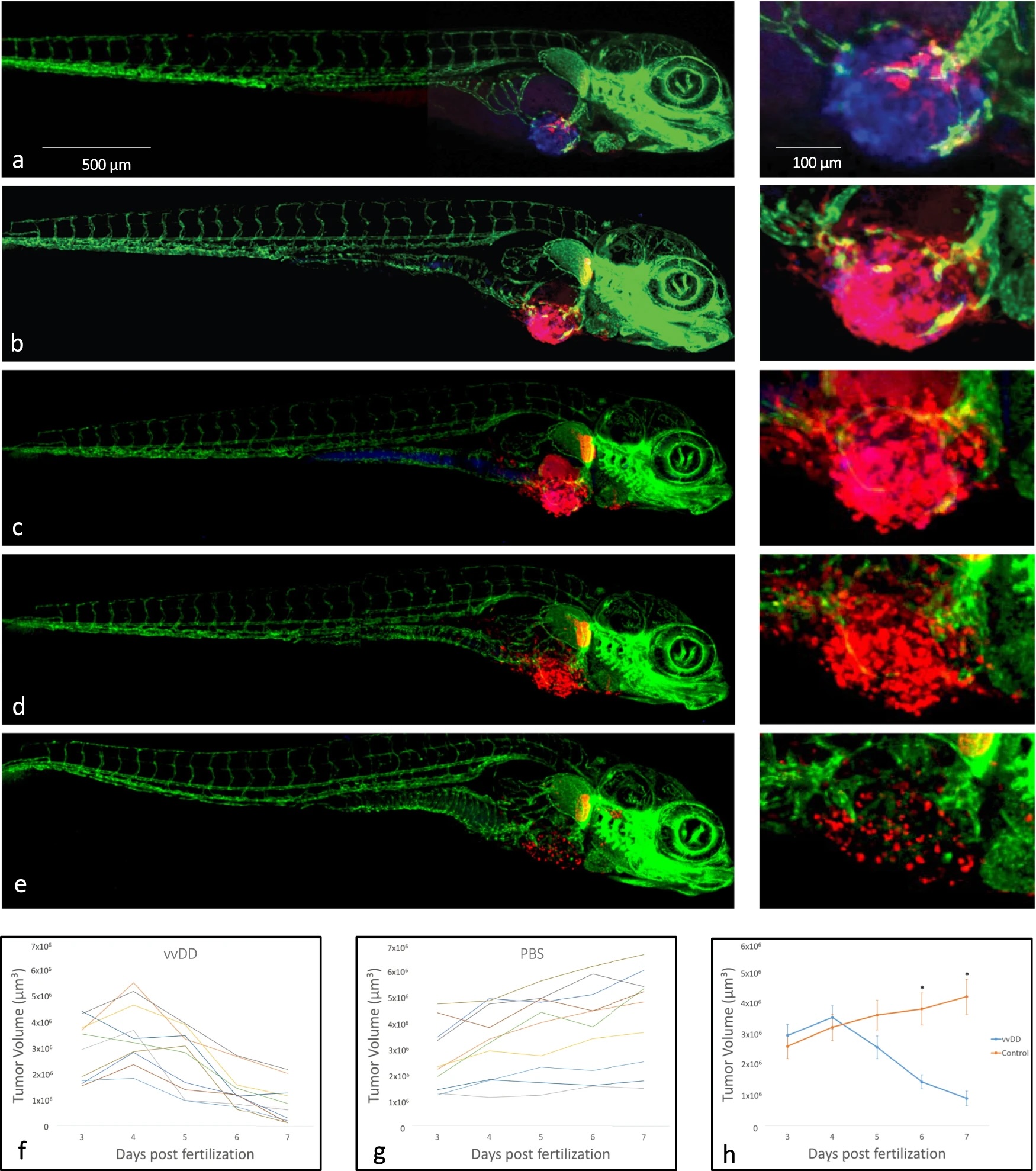

### pop_and_states.pdf

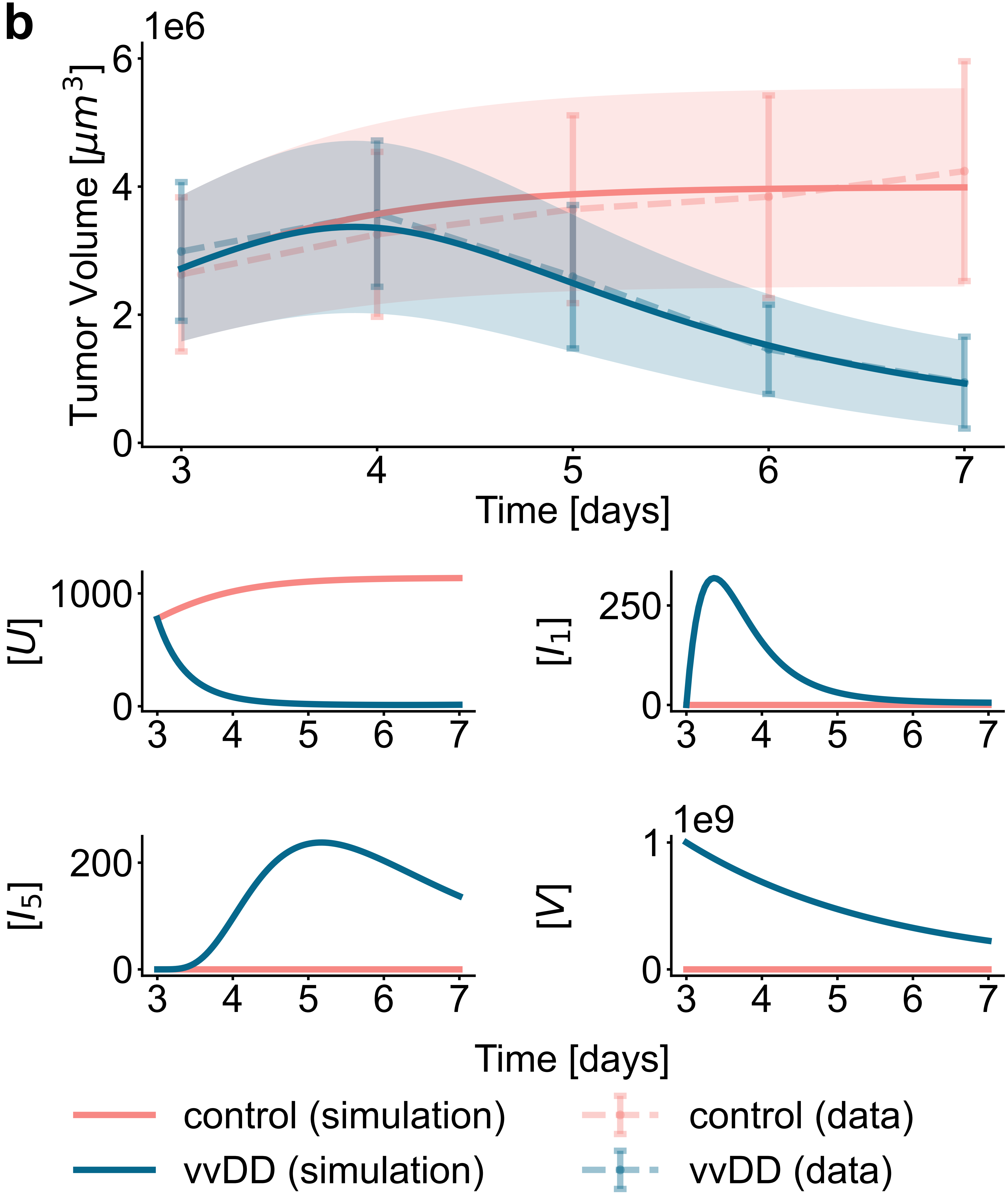

### pop_and_states.pdf

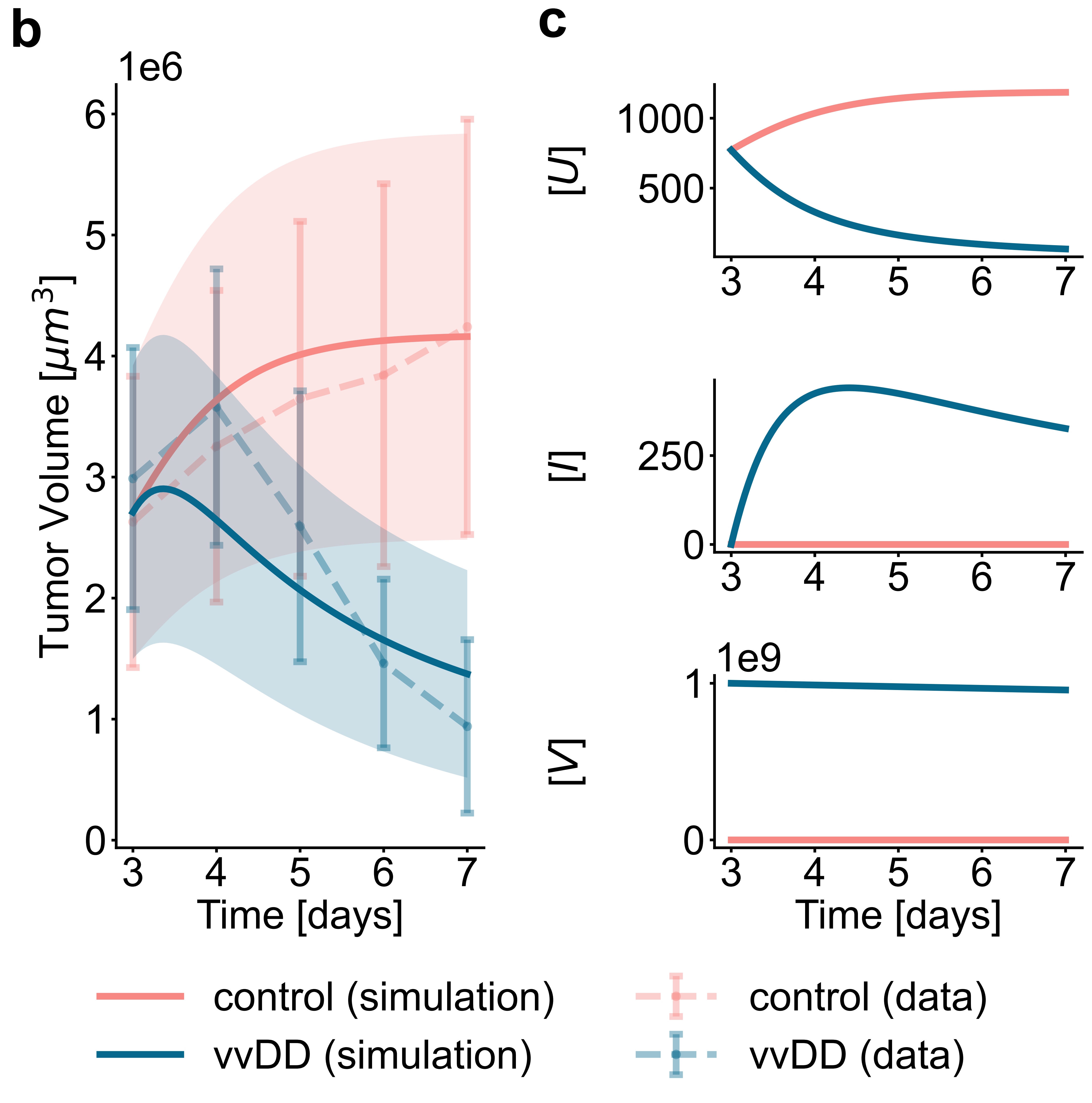

### profile_plot_res.pdf

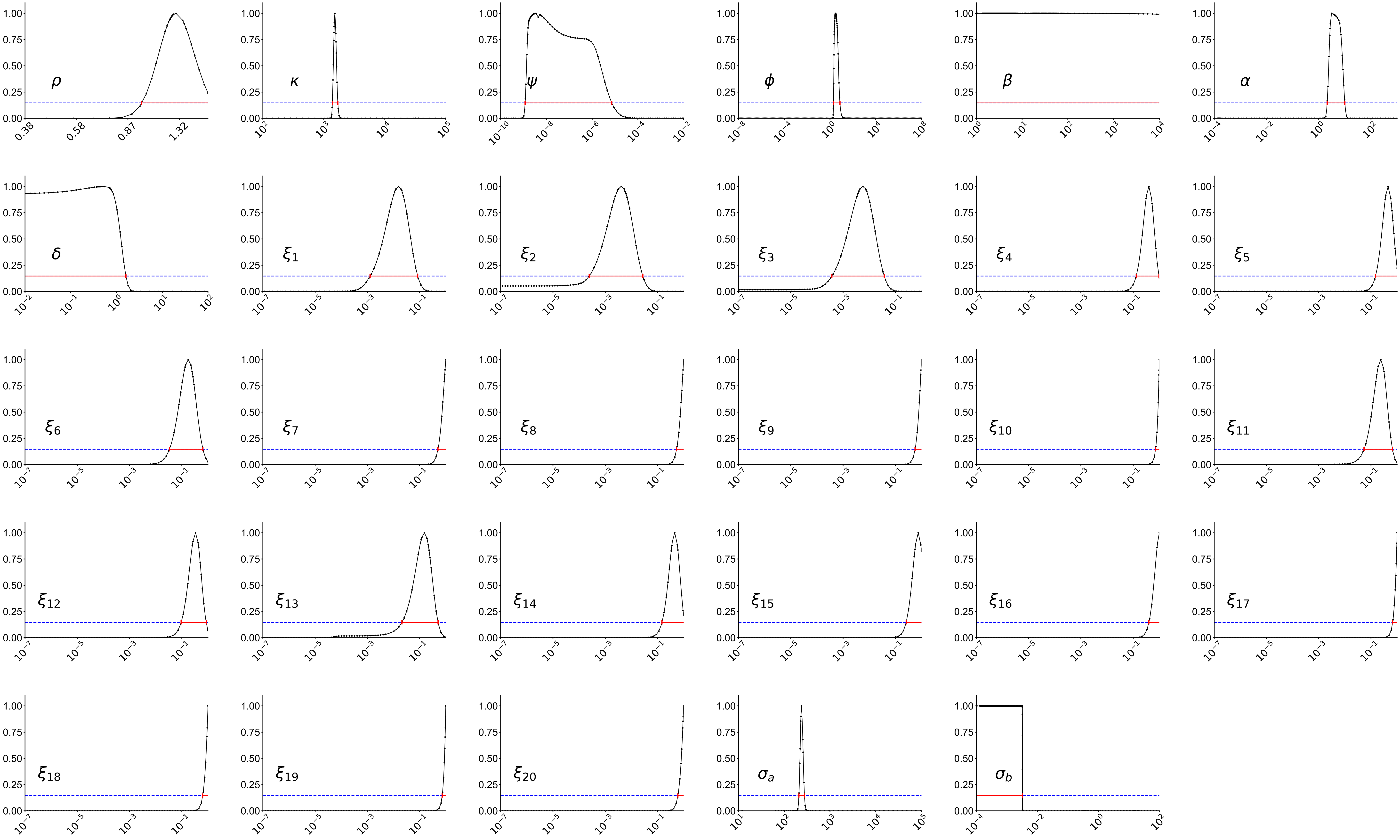

### profile_plot_res.pdf

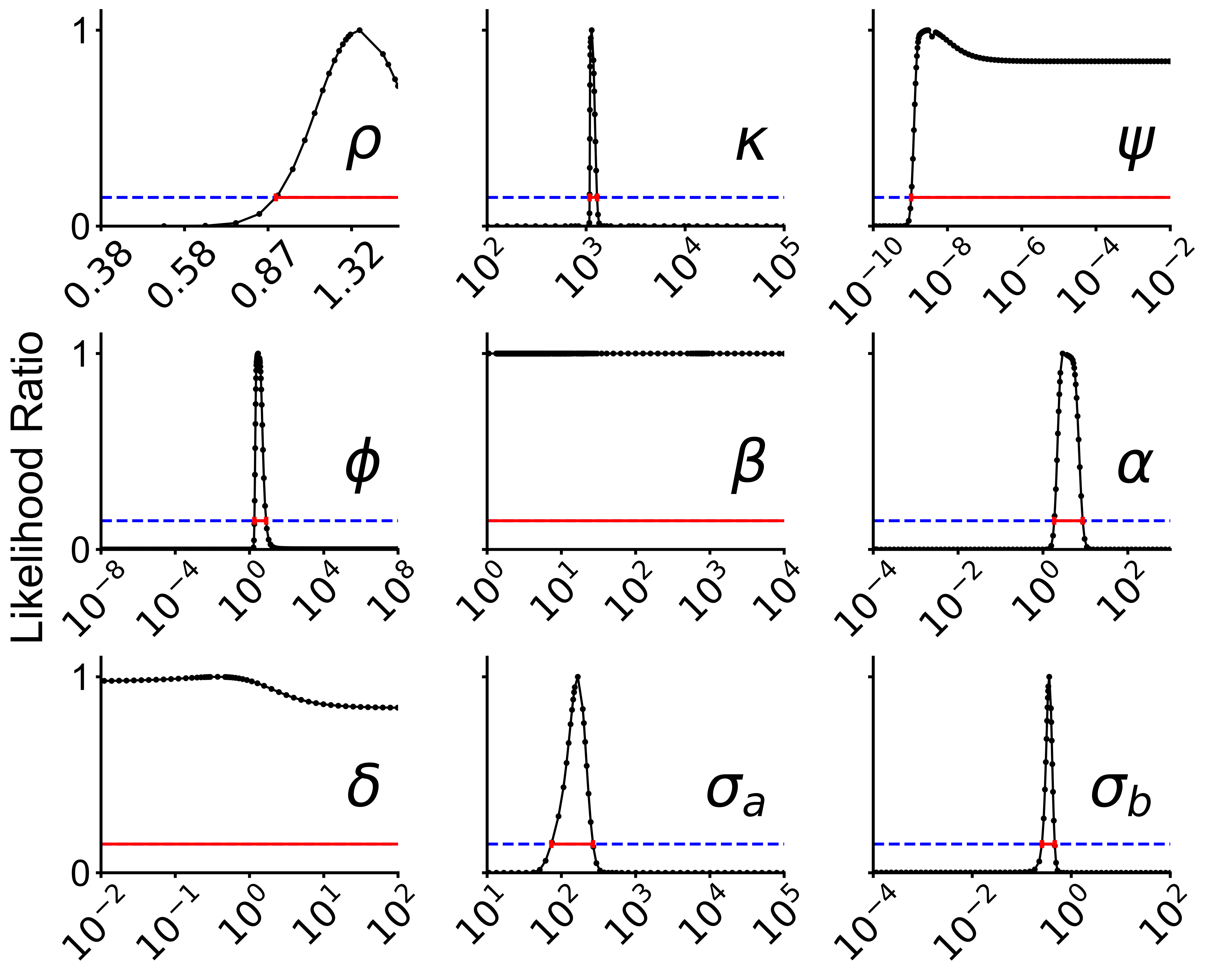

### profile_plot_res.pdf

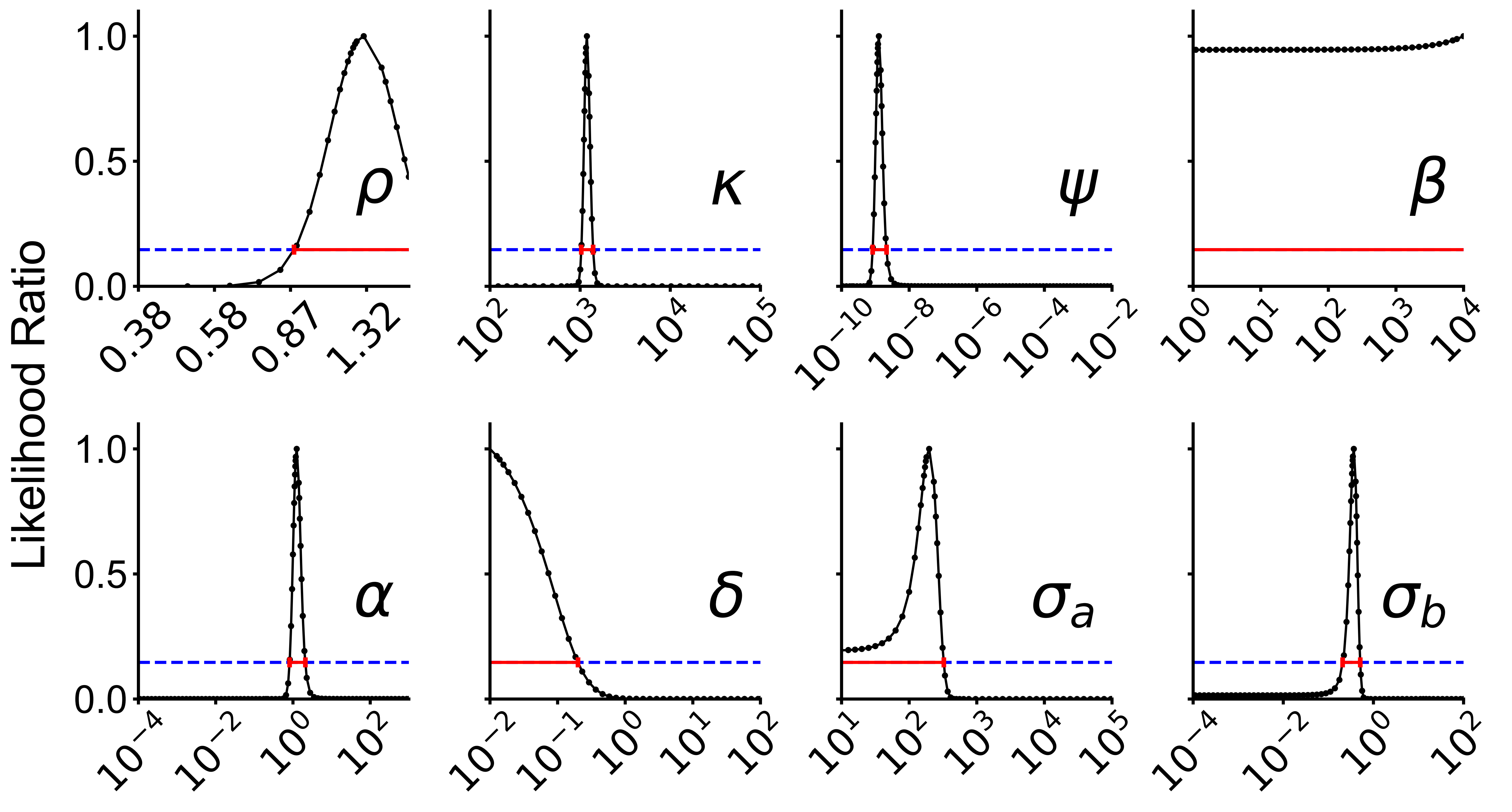

### profile_plot_res.png

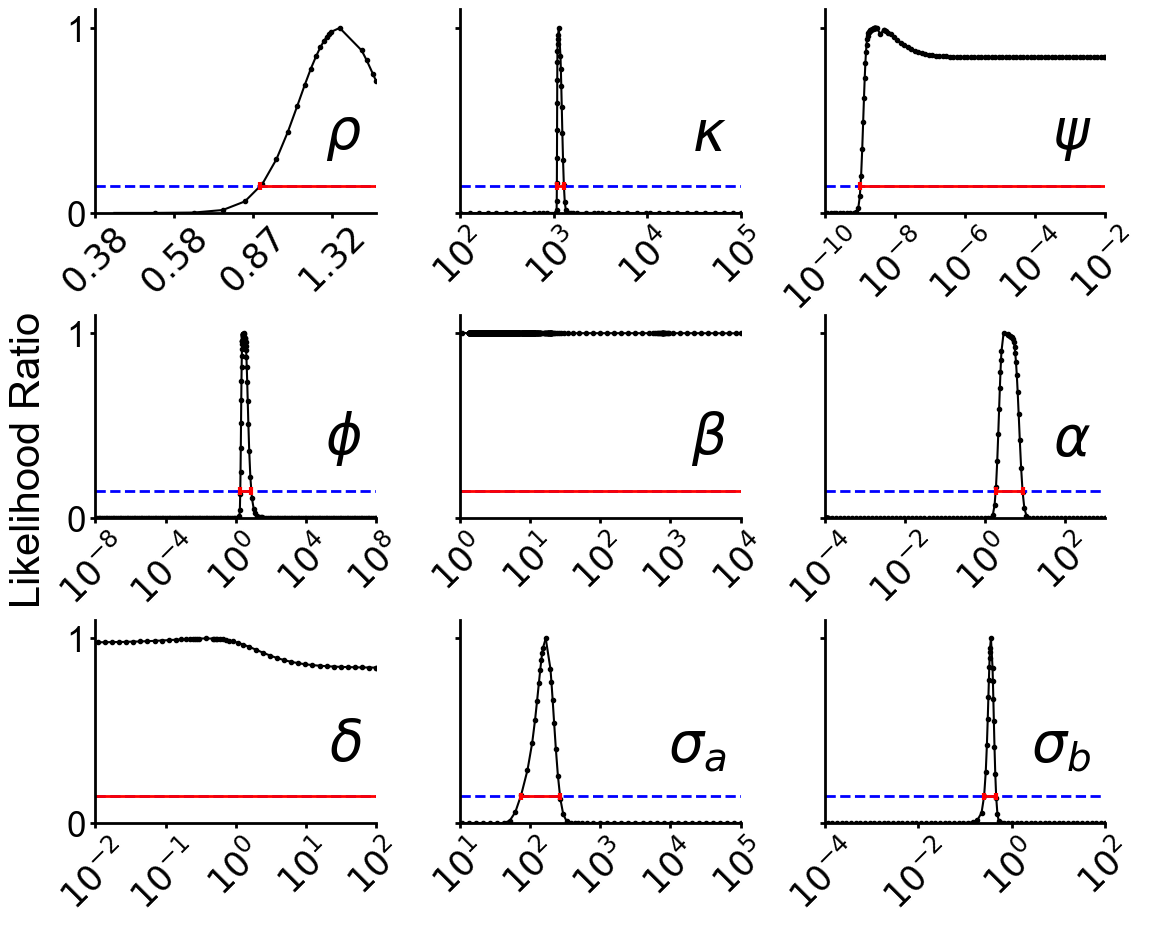

### profile_plot_res.png

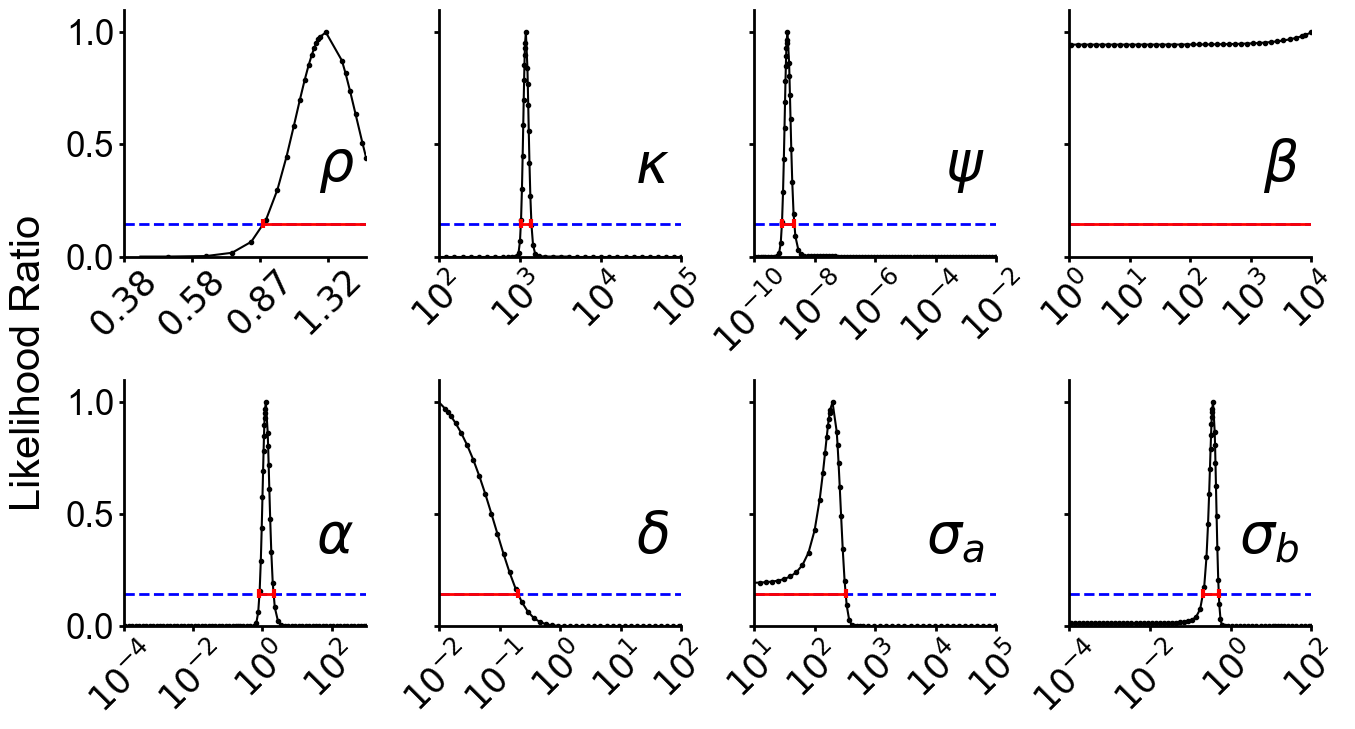

### vvDD_mechanism.png

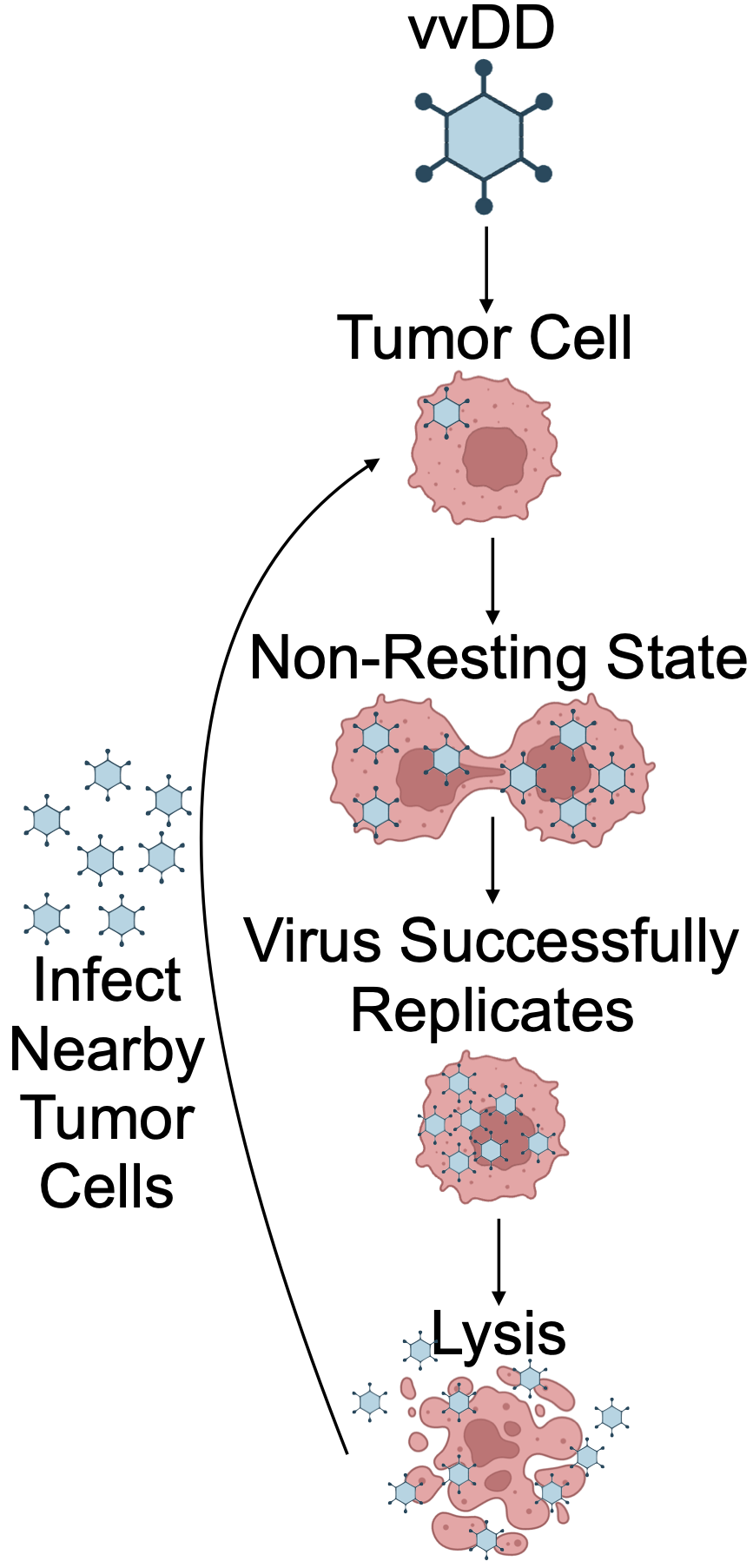

### waterfall_parameters_plot.pdf

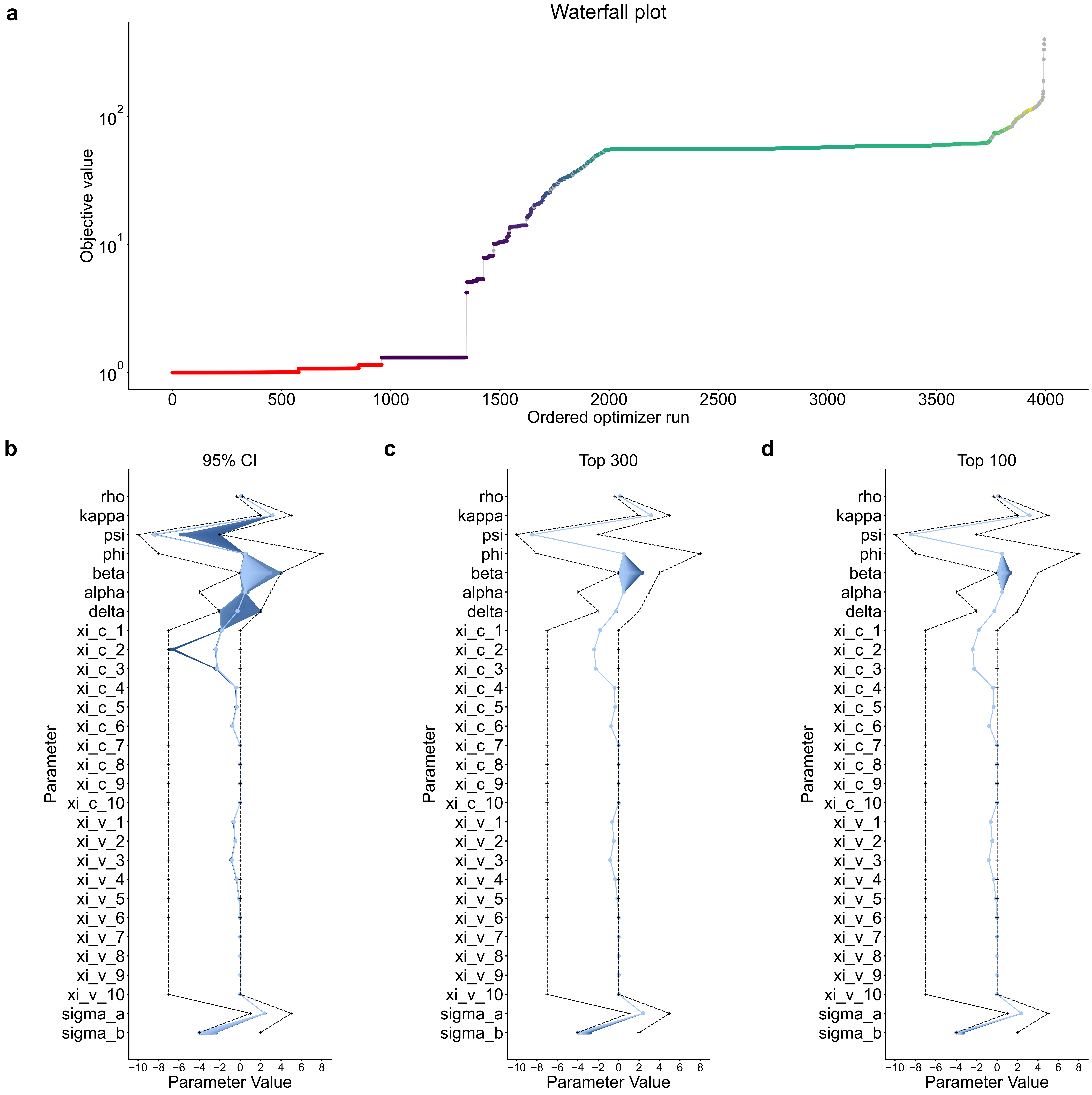

### waterfall_parameters_plot.pdf

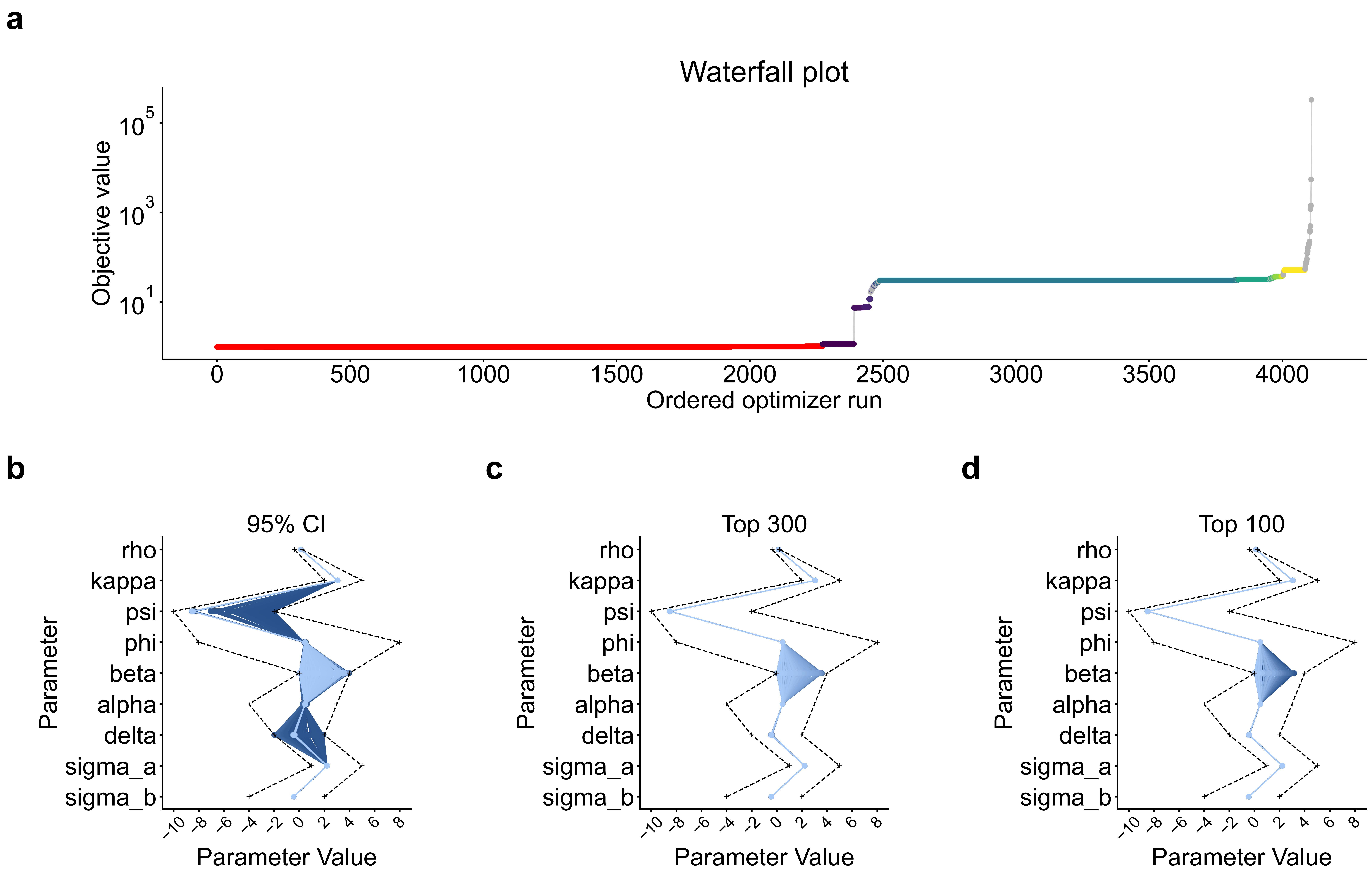

### waterfall_parameters_plot.pdf

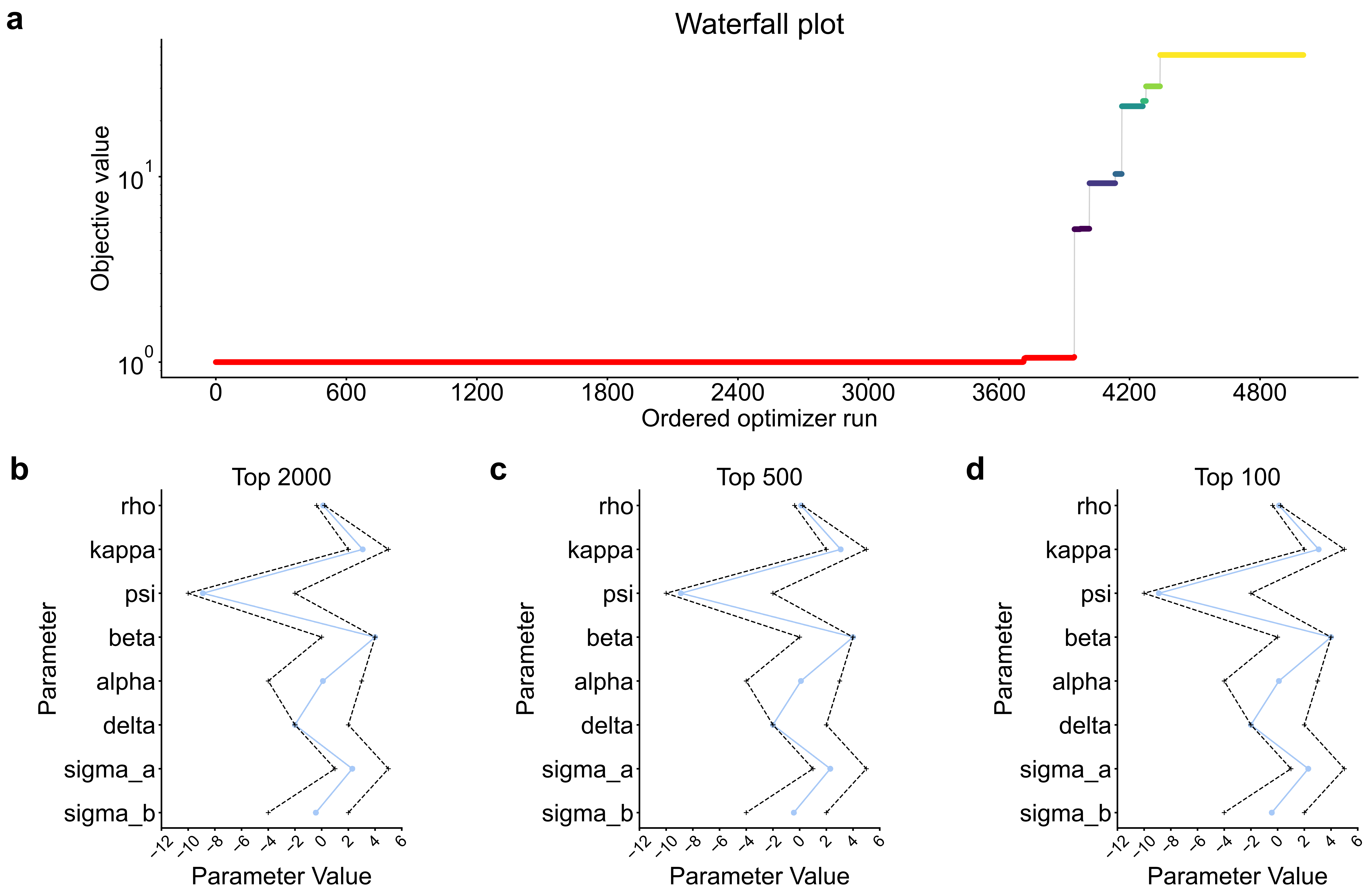
